# Mechanism of enterovirus VP0 maturation cleavage based on the structure of a stabilised assembly intermediate

**DOI:** 10.1101/2024.04.06.588229

**Authors:** Natalie J Kingston, Joseph S Snowden, Keith Grehan, Philippa K Hall, Eero V Hietanen, Tim C Passchier, Stephen J Polyak, David J Filman, James M Hogle, David J Rowlands, Nicola J Stonehouse

**Author notes:** Corresponding Authors: Natalie J. Kingston and Nicola J. Stonehouse.

## Abstract

Molecular details of genome packaging are little understood for the majority of viruses. In enteroviruses (EVs), cleavage of the structural protein VP0 into VP4 and VP2 is initiated by the incorporation of RNA into the assembling virion and is essential for infectivity. We have applied a combination of bioinformatic, molecular and structural approaches to generate the first high-resolution structure of an intermediate in the assembly pathway, termed a provirion, which contains RNA and intact VP0. We have demonstrated an essential role of VP0 E096 in VP0 cleavage independent of RNA encapsidation and generated a new model of capsid maturation, supported by bioinformatic analysis. This provides a molecular basis for RNA-dependence, where RNA induces conformational changes required for VP0 maturation, but that RNA packaging itself is not sufficient to induce maturation. These data have implications for understanding production of infectious virions and potential relevance for future vaccine and antiviral drug design.

## Introduction

Enteroviruses (EVs) comprise a large genus of positive sense RNA viruses of the family *Picornaviridae* and are jointly responsible for a range of serious diseases of both domestic livestock and humans. Examples include poliovirus (PV), human rhinoviruses (HRV) and enterovirus A71 (EVA71). EV virions are non-enveloped 30 nm icosahedral particles assembled from pentameric subunits comprising three structural proteins, VP0, VP1, VP3, surrounding the ∼7.5 kb genome. The VP0 protein is cleaved into VP2 and VP4 during the viral maturation process. This cleavage event is crucial as it both structurally stabilises and primes virus particles for subsequent infection of host cells. Mature virions attach to cells via cell surface receptors/co-receptors and the genome is released following conformational changes induced by receptor interaction and/or local environmental changes (pH and ionic changes in endosomes etc.) [1–7].

Following release into the cytoplasm of the positive sense RNA genome, translation is initiated from an internal ribosome entry site in the 5’ UTR. The open reading frame is translated as a single polyprotein which is co- and post-translationally processed into the viral structural and non-structural proteins. The viral polyprotein undergoes a series of *cis-* and *trans-* cleavage events which are mediated by two viral proteases, 2A and 3C (or the precursor 3CD). The 2A protease initiates the co-translational separation of the structural region of the polyprotein from the non-structural region [8, 9]. The 3C/3CD protease then cleaves the non-structural protein precursor into several intermediates and mature products at the VP0/3 and VP3/1 boundaries within the structural protein precursor [10–14]. The resulting structural proteins VP0, VP3 and VP1 assemble into protomers 5 assemble into pentamers, twelve of which assemble into icosahedral particles in the presence of a nascent VPg-coupled genome [15–19] (Fig 1). These transient intermediate particles have been termed provirions, comprising the full complement of structural proteins encapsidating the RNA genome. A series of conformational changes within the structural proteins ultimately stabilise the particle and result in the processing of VP0 into VP4 and VP2. In the absence of genome packaging, empty capsids (ECs) form, but these particles do not process the VP0 precursor and are less stable (Fig 1). When retained in their native conformation, these ECs can reversibly dis- and re-associate into icosahedral particles, but readily undergo an irreversible expansion and this ability for reversible disassembly is lost [19].

**Figure 1:**
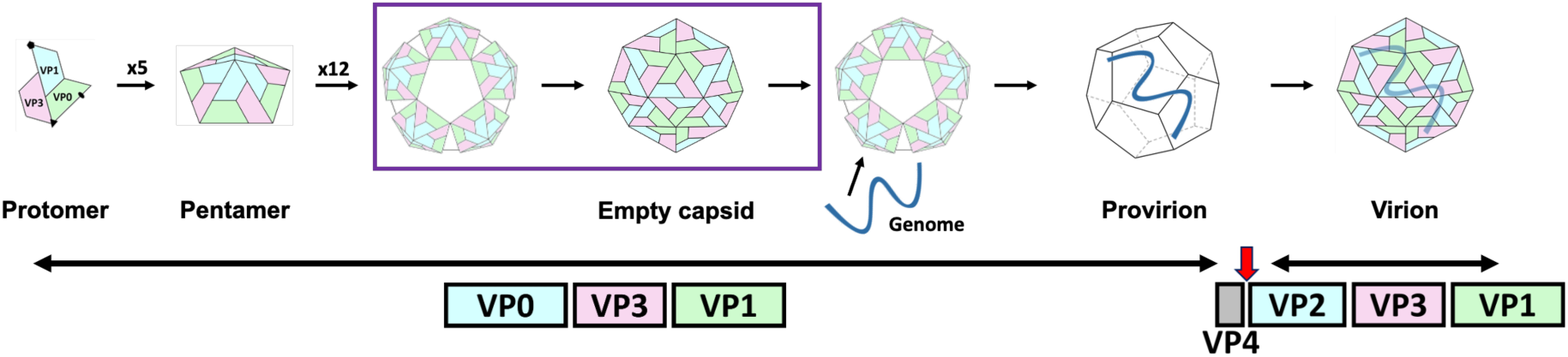
Schematic of enterovirus assembly. Five protomers of VP0, VP3 and VP1 assemble into a pentamer after the P1 precursor is cleaved by 3C or 3CD at the VP0/3 and VP3/1 boundaries. Empty capsids form when 12 pentamers assemble in the absence of genome, these may be an off-pathway product or may partially or fully disassemble before reassembly with the RNA genome (indicated by purple box). Provirions exist briefly before genome packaging and the cleavage of VP0 into VP4 and VP2 (red arrow). VP4 is internal, thus is not depicted as a part of the capsid.

The cleavage of VP0 occurs rapidly upon encapsidation of the viral genome, consequently, assembly intermediates (provirions) are challenging to detect and investigate. However, by using sensitive methods such as radio-labelling, provirions can be detected in wild type (WT) PV infected cellular extracts [20, 21]. Furthermore, mutations which inhibit VP0 cleavage have been shown to result in the accumulation of provirions [22, 23]. Indeed, Compton *et al*. (1990) generated a mutant of PV (VP2 R76Q) which showed a defect in VP0 cleavage when grown at 39°C but had a near WT phenotype when cultured at 32°C [23]. However, it is unclear how this mutation results in a cleavage-defective phenotype as the altered residue is located on the outer surface of the capsid.

The rapid rate of particle maturation of WT EVs has made it particularly challenging to define a catalytic mechanism or to understand the role of RNA packaging in particle maturation. The historic basis for the currently accepted model of VP0 maturation cleavage is primarily based upon the structure of an unexpanded (native) PV EC (PDB: 1pov). This structure revealed several interesting conformational features and ultimately led to the widely accepted model for VP0 maturation cleavage in EVs. The initial work by Basavappa *et al*. [24] using PV showed the scissile boundary interacting with a highly conserved histidine residue which was proposed to provide a catalytic function. In this conformation, the serine residue in the P1’ position immediately downstream of the cleavage site interacts with the putative catalytic histidine residue (VP2 H195) [24]. This interaction requires that the main chain residues flanking the scissile site adopt an unfavourable combination of φ and ψ torsions and results in the carbonyl oxygens of the P1 and P1’ residues being oriented together to form a negatively charged pocket. The combination of unusual torsions surrounding the scissile site and the polarisation of the carbonyl groups would be expected to destabilise the scissile peptide bond suggesting that the scissile boundary is appropriately primed for VP0 cleavage. In their model, Basavappa *et al*. proposed that interactions of a cation or a base from the packaged RNA with the pair of carbonyl oxygens would hyperpolarise these carbonyl groups, ultimately leading to nucleophilic attack and VP0 cleavage [24]. Additionally, while Basavappa *et al*. predicted that an RNA hairpin would occupy the a trefoil-shaped pocket formed by the three internal clefts at the 3-fold region with the tip of the hairpin approaching the scissile bond, RNA has not been directly detected in this region [24]. In mature EV virions, RNA has been directly detected interacting with a well-conserved tryptophan residue (VP0 W107) [25–27], and the downstream tyrosine residue (VP0 Y110) [25].

Hindeyah *et al*. demonstrated the functional importance of the key histidine by showing that mutation of this residue resulted in PV particles which packaged genome but did not cleave VP0 [22]. These provirions were presumed to represent stabilised assembly intermediate particles unable to complete VP0 cleavage, thus confirming the involvement of VP0 H195 in the maturation process, possibly by “activating” an adjacent water molecule which would then serve as the nucleophile in the hydrolysis of the activated peptide bond.

High-resolution structures of PV1 and human rhinovirus C (HRVC) unexpanded ECs (PDB: 1pov and 5jzg, respectively) and mature virions (PDB: 1hxs and 5k0u, respectively) provide details of protein conformations at the start and end of the maturation process [24, 28, 29]. However, until now there has been no structural information on the conformational details of the intermediate provirion particles. Such structures are invaluable for further understanding the details surrounding the conformational changes which lead to particle maturation.

Recent improvements in both cryoEM and molecular biology approaches for the large-scale recovery of non-infectious virus have allowed us to make significant progress in understanding these intermediate-state provirions. We have generated a mutant virus of EVA71 which contains a full complement of structural proteins and the viral genome but fails to cleave VP0. This allowed us to determine the first high-resolution structure of a picornavirus provirion and provide detailed information on the intermediate conformations required for VP0 cleavage.

This structure confirms the importance of the conformation described in Basavappa *et al.* [24], and suggests a molecular mechanism by which RNA packaging disrupts stabilising interactions, resulting in conformational changes and VP0 cleavage and indicating at least one direct role for RNA in the initiation of VP0 cleavage in EVs. Finally, this structure identifies critical residues required for particle maturation and provides sufficient detail to present a model for picornavirus VP0 maturation which is well supported by bioinformatic data.

## Methods

### Bioinformatic analysis and mutant selection

The published nucleotide sequence of all EVs with genome lengths between 7000-8000 nt were imported into Geneious Prime 2023.0.4 from GenBank. The polyprotein encoding region of the genome was translated and a pairwise sequence alignment was performed using MAFFT [30] on Geneious Prime. The alignment, containing 7955 sequences, was manually inspected for regions of high sequence conservation, regions of interest were subsequently mapped to published EV structures at different states of assembly (native and expanded states of virus and EC). Several regions of interest were identified based upon sequence conservation and conformational position. The residue VP0 E096 was selected for further investigation, several other residues of interest were identified but are not described here (Fig S1).

### Recovery of EVA71

Wild type or mutant EVA71 viruses were recovered from *in vitro* transcribed RNAs, as previously described [31]. Briefly, WT, mutant or replicon encoding plasmids were linearised and purified by phenol/chloroform extraction. RNA was synthesised using the RiboMaX T7 express large-scale RNA production system (Promega). Samples were purified through an RNA clean and concentrator column (Zymo research, USA), before electroporation into HeLa cells within a 4 mm cuvette, using a square wave at 260V for 25 milliseconds. Cells were incubated at 37°C with 5% CO_2_ in a humidified chamber until harvest. Samples used for molecular characterisation were harvested at 18 hours post-electroporation from the supernatant, samples generated for structural studies were harvested 8 hours post-electroporation and isolated directly from cells.

### RTqPCR

Samples collected from sucrose gradients were diluted 10-fold in nuclease-free water supplemented with RNA-secure. Samples were heated to 60°C for 10 minutes to both activate the RNA-secure and expand particles to allow genome egress. Samples were then used in a one-step RTqPCR (Promega) per manufacturer’s instructions. Samples were assessed alongside a titrated virus sample purified in the same manner to generate a standard curve. Total genome copies were determined from this standard curve, and data are displayed as percentage genome contained within each fraction.

### Virus purification

Virus samples were harvested from cells after 18 hours by addition of NP40 to a final concentration of 0.02%, agitating briefly and subjecting to a single cycle of freeze-thaw to facilitate release of virus particles from cells. The contents of wells were collected and clarified at 17,000 rcf for 10 minutes. The clarified supernatants were then used directly for sucrose gradient purification and serial passage. Samples were loaded on top of discontinuous 15-45% sucrose gradients and centrifuged at 50,000 rcf for 12 hours. Seventeen 1 ml fractions were collected (top-down) and assessed for the presence of VP0 and VP2 by western blot with mAb979 (Merck Millipore).

### Virus purification for cryoEM

Cells collected 8 hours after electroporation were resuspended in lysis buffer (PBS supplemented with 1% v/v NP40 and 0.5% w/v sodium deoxycholate) and incubated with agitation for 2 hours at 4°C. Samples were supplemented with MgCl_2_ to a final concentration of 1.5 mM before the addition of 25 U/ml Denerase (c-Lecta). Samples were incubated overnight at 4°C before clarification at 4,000 rcf for 30 minutes. Clarified supernatants were pelleted through 30% sucrose cushions at 150,000 rcf for 3.5 hours and re-suspended in 1 ml PBS. Samples were loaded on top of a 15-45% sucrose gradients and separated, as above. Peak fractions were subsequently diluted 1:1 in PBS and underlaid with a 20-45% discontinuous sucrose gradient before centrifugation at 150,000 rcf for 3 hours. Fractions were collected (as above) and assessed for the presence of VP0 by western blot. Peak fractions were concentrated across a 100 kDa mwco spin concentrator, excess sucrose was removed with successive rounds of dilution in PBS and concentration (as previously described)[31].

### Electron microscopy

CryoEM grids were prepared, data collected and processed as previously described [32], with some small modifications. Briefly, 3 µl of concentrated sample was applied to ultrathin continuous carbon-coated lacey carbon 400-mesh copper grids (Agar Scientific, UK) following glow discharge in air (10 mA, 30 s). Sample application was performed in a humidity-controlled chamber maintained at 8°C and 80% relative humidity. Excess liquid was removed by blotting either immediately or following a 30-s incubation period, using a range of blotting times (1.5 s to 3.5 s). Grids were vitrified in liquid nitrogen-cooled liquid ethane using a LEICA EM GP plunge freezing device (Leica Microsystems, Germany). Screening and data collection were performed on an FEI Titan Krios transmission EM (ABSL, University of Leeds) operating at 300 kV. Imaging was performed at a magnification of 130,000× and with a calibrated object sampling of 0.91 Å/pixel (a complete set of data collection parameters are provided in Table S1).

### Image processing

Image processing was performed using Relion 3.1.1 [33, 34]. Motion-induced blurring was corrected using the Relion implementation of MotionCor2 [35], then CTFFIND-4.1 was used to estimate CTF parameters for each micrograph [36]. Particles were picked and extracted and underwent 2D classification; those from selected 2D classes were re-extracted and subjected to 3D classification into four classes. The particles in each class were then processed separately, through 3D refinement using *de novo* initial models (with icosahedral symmetry imposed), CTF refinement and Bayesian polishing. A final 3D refinement was performed with a mask to exclude solvent and calculation of solvent-flattened Fourier shell correlations (FSCs). Sharpening was performed using a solvent-excluding mask, and the nominal resolution was calculated using the ‘gold standard’ FSC criterion (FSC=0.143) [37]. Local resolution was determined using Relion.

To investigate the internal capsid density a focussed classification approach was employed, as previously described [38–42]. Briefly, particles contributing to the final reconstruction were symmetry expanded, using the relion_symmetry_expand tool. A cylindrical mask was created in SPIDER [43], and the mask placed over the capsid features of interest before being subjected to 3D classification. Full reconstructions were generated from the classes without symmetry imposed using the relion_reconstruct tool.

### Model building and refinement

To build an atomic model for the particles, an atomic model of a native EVA71 virion (PDB: 3vbs) was rigid-body fitted into a single asymmetric unit within the density map using UCSF Chimera [44]. The model was altered to represent the provirion sequence in Coot and regions of the peptide backbone without supporting density were removed. Iterative cycles of inspection and manual refinement in Coot [45], followed by automated refinement in Phenix [46], were performed to improve atomic geometry and the fit of the model within the density. To avoid erroneous positioning of the model within density corresponding to adjacent protomers, density from adjacent asymmetric units was occupied with symmetry mates during automated refinement. Molprobity was used for model validation [47].

### Structure analysis and visualisation

Visualisation of structural data was performed in UCSF Chimera [44], UCSF ChimeraX [48] and PyMOL (The PyMOL Molecular Graphics System, Version 2.1, Schrödinger, LLC).

Atomic models used for reference throughout this work are described in Table 1.

**Table 1:** Atomic models used as references.

| PDB | Virus | State | Particle | RNA detected | Reference |
| --- | --- | --- | --- | --- | --- |
| 3vbs | Enterovirus A71 | native | virion |  | [49] |
| 3vbu | Enterovirus A71 | expanded | empty capsid |  | [49] |
| 1hxs | Poliovirus 1 | native | virion |  | [29] |
| 1pov | Poliovirus 1 | native | empty capsid |  | [24] |
| 5jzg | Human Rhinovirus C | native | empty capsid |  | [28] |
| 5k0u | Human Rhinovirus C | native | virion |  | [28] |
| 2plv | Poliovirus 1 | native | virion | Yes | [25] |
| 5c9a | Coxsackievirus A16 | native | empty capsid |  | [50] |
| 5c4w | Coxsackievirus A16 | native | virion |  | [50] |
| 5c8c | Coxsackievirus A16 | native | virus-like particle |  | [50] |
| 6thn | Bovine enterovirus | native | virion | Yes | [26] |
| 7bg6 | Human Rhinovirus 14 | native | virion | Yes | [27] |

### Picornavirus bioinformatic analysis

Picornavirus sequences were retrieved from GenBank for 62 species. Two species, *Fipivirus* and *Pemapivirus*, were excluded from the analysis due to unavailability of suitable sequence data. Collected sequences were filtered to further exclude synthetic, unverified, and patent sequences. Sequence data were split into species specific subsets, further categorized by their current ICTV assigned VP0 cleaving (N=16240) and non-cleaving (N=1694) classification. Coding sequences (CDS) were extracted based on the sequence annotation. In cases where annotation was lacking for part of the sequences, a species-specific alignment was created, and annotations transferred to determine the CDS. All multiple sequence alignments (MSA) were carried out using MUSCLE v5.1 [51] in Geneious Prime 2023.2.1 using the default settings.

P1 regions were similarly extracted based on the sequence annotations. In cases where the species data completely lacked annotations to determine the P1 region, neighbour-joining trees were built for full polyprotein data and closest related species to the query species were used as a reference to determine the P1 region through a multiple sequence alignment. Additionally, if established features such as the 2A protein N/HC-box or the NPGP motif were present, these were further used to establish the P1 region. The region of interest (ROI) for the VP0 cleaving viruses was extracted from the P1 region multiple sequence alignments based on the conserved features (conserved L on the N-terminus, conserved H on the C-terminus). For the VP0 non-cleaving viruses, the full VP0 region was extracted instead.

From the above species specific MSAs for each region (full polyprotein, P1, ROI/VP0), majority consensus sequences were extracted. Extracted consensus sequences for each species within each sequence set were combined with an infectious flacherie virus (IFV) genome (acc. No. NC_003781) (a picornavirus from another family) to act as an outgroup. These consensus sequence MSAs constructed for each genomic region were stripped of gaps (90% threshold per site), and alignments were inspected and fixed for notable major issues such as large portions of missing data.

Phylogenetic analyses were carried out with the MSAs of the species-specific consensus sequences using IQTree2 [52]. IQTree2 analysis used the built-in ModelFinder to find the best fit evolutionary model for each MSA, with subsequent maximum-likelihood inference carried out together with ultra-fast bootstrapping. Bootstrapping analysis was conducted using enough replicates until convergence was achieved through the SH-aLRT test. Consensus trees for each MSA as built by IQTree2 were exported in Newick format and analysed in R. Tanglegram analysis of the constructed phylogenies was carried out using a custom script and the dendextend v1.17.1 [53] and phylogram v2.1.0 [54] packages. Tanglegrams were constructed comparing different regions (full polyprotein, P1, VP0/ROI) within each sequence data set against each other to highlight possible topological incongruence between the trees.

## Results

### Identification of conserved residues within the Enterovirus genus

Many regions of the polyprotein are highly conserved across EVs and several of these features are associated with known functions. For example, a H-E-C catalytic triad of 3C^pro^, Q-G at 3C^pro^ cleavage boundaries, G-D-D at the active site of the RNA-dependent RNA polymerase (3D^pol^) and a P-H-Q motif in VP0, of which the histidine is important for VP0 maturation cleavage [12, 24, 55–58]. Given the functional importance of these highly conserved residues, we investigated whether critical functional roles can be associated with other conserved residues.

A broad analysis of all the available EV protein sequences was performed to identify highly conserved features and investigate the potential roles they may have in RNA-packaging and the induction of VP0 cleavage. Several candidate residues were identified, with VP0 096 as the focus of the work described here. This residue is conserved as a negatively charged amino acid (Asp or Glu) in >99.9% of all EV sequences (Fig S1) and was of particular interest as the VP0 A_2_B-loop, where E096 resides, is flexible and adopts multiple conformations across different assembly states.

### Generation of mutationally stabilised E096A EVA71 provirion

To assess the phenotypic consequences of mutating this residue, an infectious clone with a E096A mutation in EVA71 was generated and investigated alongside a WT control. *In vitro* transcribed RNAs derived from the control or mutated sequences were electroporated into susceptible cells to recover the cognate viruses. The presence of assembled viral particles in extracts from the electroporated cells was determined by sucrose gradient ultracentrifugation. As expected, the WT virus separated into two distinct populations, the more slowly sedimenting particles contain unprocessed VP0, consistent with ECs, and a faster sedimenting peak of particles containing VP2, consistent with mature virions (previously described [31]) (Fig 2). Particles generated by the E096A mutant virus contained no detectable VP2, suggesting a defect in the cleavage of VP0. Although the majority of particles sedimented similarly to ECs, some were detected in the fractions between virions and ECs, consistent with the previously described properties of PV provirions [22, 23]. The presence of genome within these particles was demonstrated by RTqPCR, suggesting that the VP0 E096A mutant can encapsidate viral RNA despite the absence of detectable VP0 cleavage (Fig 2). As expected for the WT virus, sample genome was maximally detected in fractions which contained VP2, corresponding to mature virions (Fig 2).

**Figure 2:**
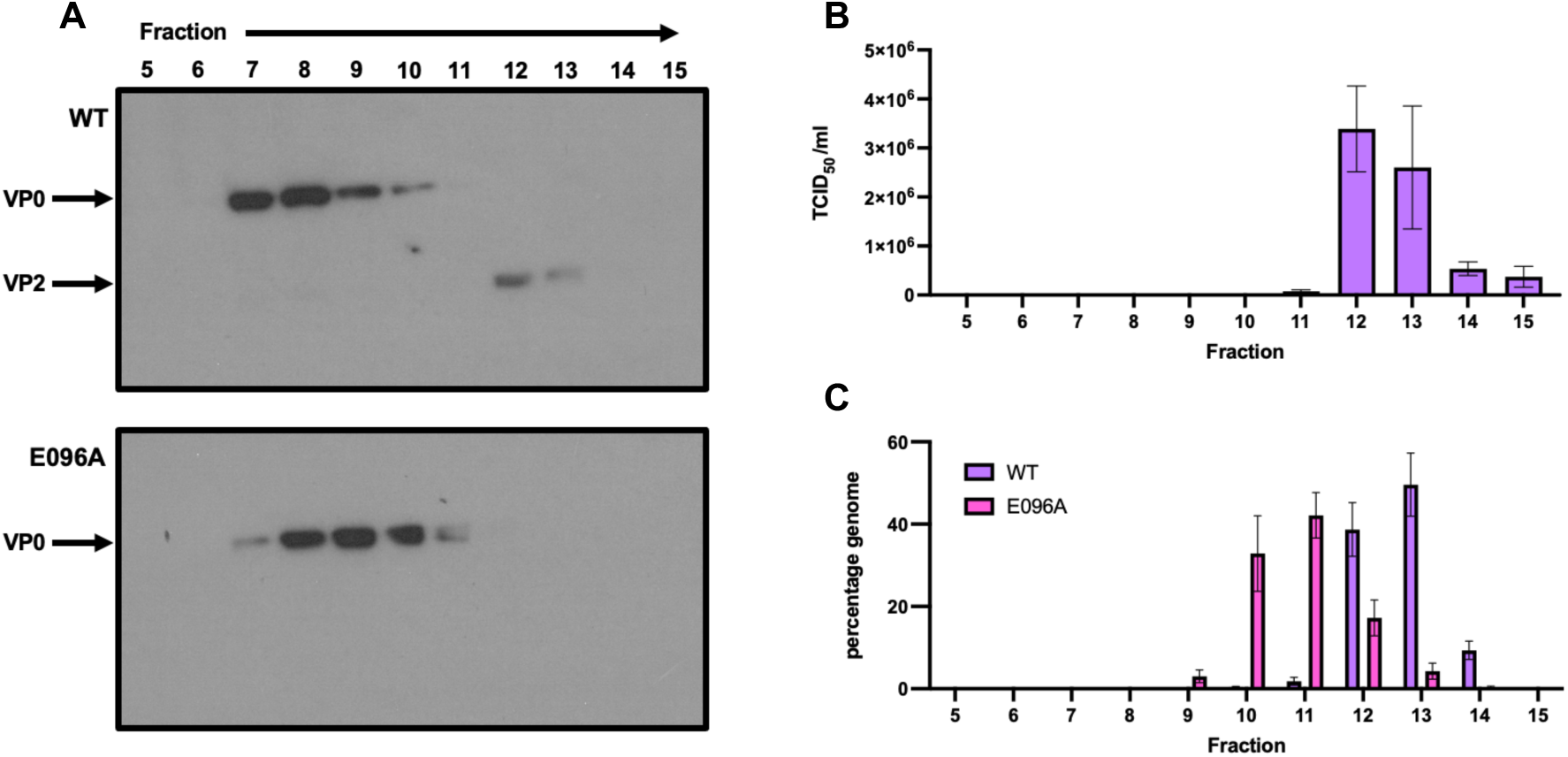
Molecular characterisation of EVA71 provirions. **A**) Samples of WT and E096A mutant EVA71 were recovered from *in vitro* transcribed RNA electroporated into HeLa cells and separated on 15-45% sucrose gradients. Samples were collected top-down and fractions 5-15 of 17 analysed by western blot using mAb979, which recognises both VP0 and VP2. **B**) Infectious virus titre of WT virus fractions presented as TCID_50_/ml, graphed mean ± SEM. **C**) The presence of genomic RNA in these fractions was assessed by RTqPCR and was quantified relative to a titrated viral sample produced in the same manner. Genome content is presented as percentage genome and is graphed mean ± SEM.

Together these data indicate that the E096A mutation was compatible with virus assembly, including genome packaging, but not with VP0 cleavage. Infectious virus was not derived from the E096A mutant virus despite serial passage, emphasising the requirement for VP0 cleavage in the generation of infectious particles (Fig S2).

To produce sufficient material for structural characterisation, the E096A mutant virus was recovered directly from cells transfected with *in vitro* transcribed RNA. Samples were purified *via* sequential sucrose gradients before being concentrated using 100 kDa mwco spin concentrators. Samples were visualised by negative stain TEM, which revealed both rounded (native) and angular (expanded) particles, consistent with incomplete separation of ECs and provirions (Fig S3).

### Structural analysis of E096A provirions by cryoEM

The provirion containing fractions were analysed by cryoEM to better understand the role of E096 in the prevention of viral maturation cleavage. A total of 8964 particles were collected of sufficient quality to contribute to a reconstruction. These particles underwent 2D and 3D classification.

Visual inspection of negative stain TEM micrographs revealed at least two distinct particle morphologies within the sample (Fig S3). Incomplete separation of genome-containing and empty particles along with the potential for particle expansion suggested several possible conformations may be present within the sample. We thus elected to separate particles into four 3D classes. The class containing the majority of particles (4897 particles, ∼55%) was resolved to 3.4 Å. The morphology of this class was consistent with an expanded state and was not analysed further. A small population (146 particles) did not contribute to a reconstruction, but the remaining 3921 particles appeared to be of a native conformation and occupied two classes.

Initial maps showed one class containing 2739 particles (∼31%) which were likely to be native conformation ECs, with no evidence of internal density corresponding to genome. The other class contained 1182 particles (∼13%), and these showed internal density reaching to the shell of the capsid, consistent with genome-containing particles (provirions) (Fig 3). After several cycles of refinement (3D auto-refinement, post processing, CTF refinement, and Bayesian polishing) the native ECs were resolved to 2.5 Å and the genome-containing provirion resolved to 2.7 Å using the gold-standard FSC (0.143) (Fig 4, Fig S4).

**Figure 3:**
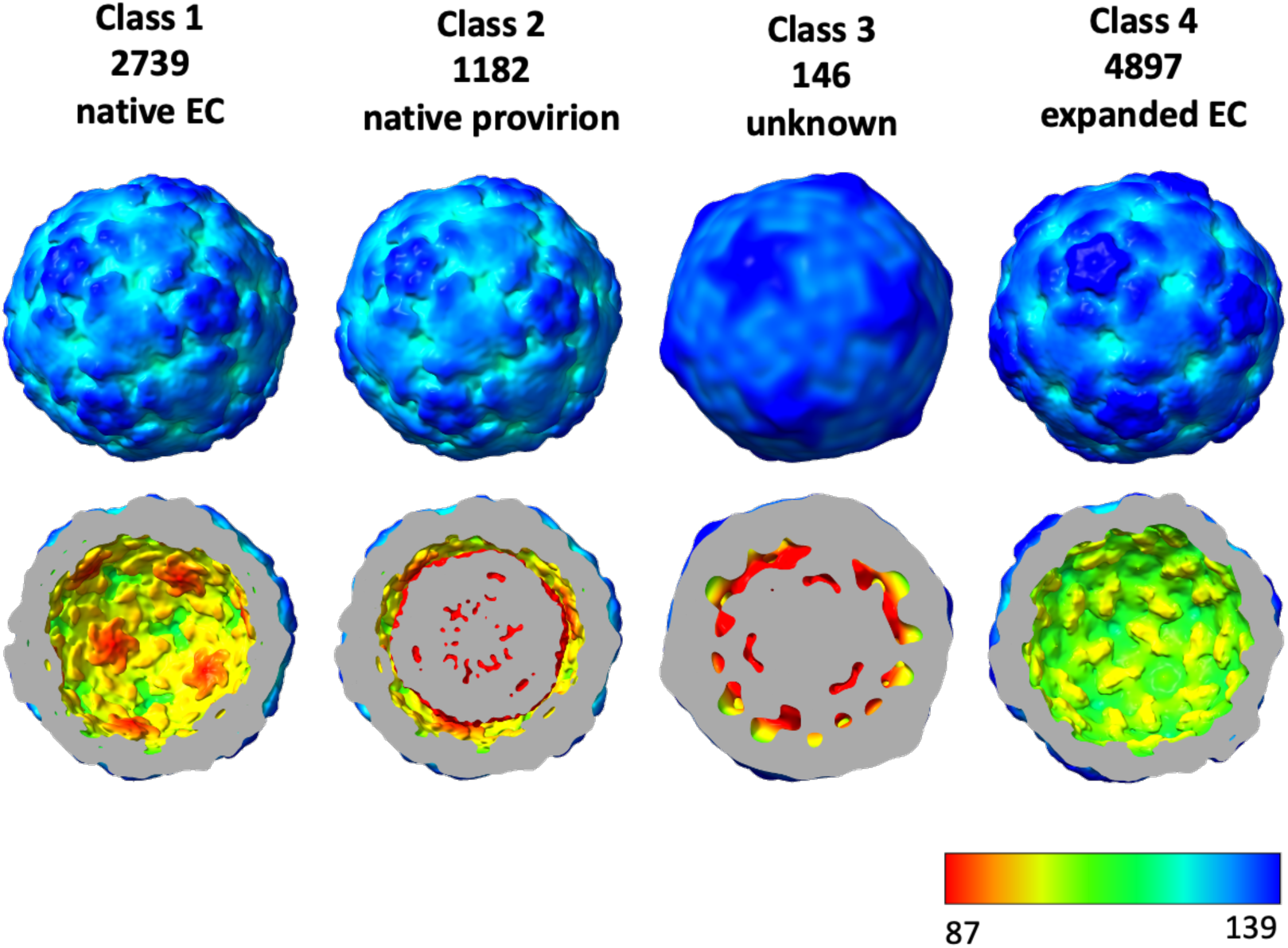
Isosurface representation of EVA71 E096A particle classes: Complete and sectional isosurface representation of EVA71 E096A particle classes. Particles coloured by radial distance in Å, indicated in the bottom key. The clipped surface of the particles is coloured grey.

**Figure 4:**
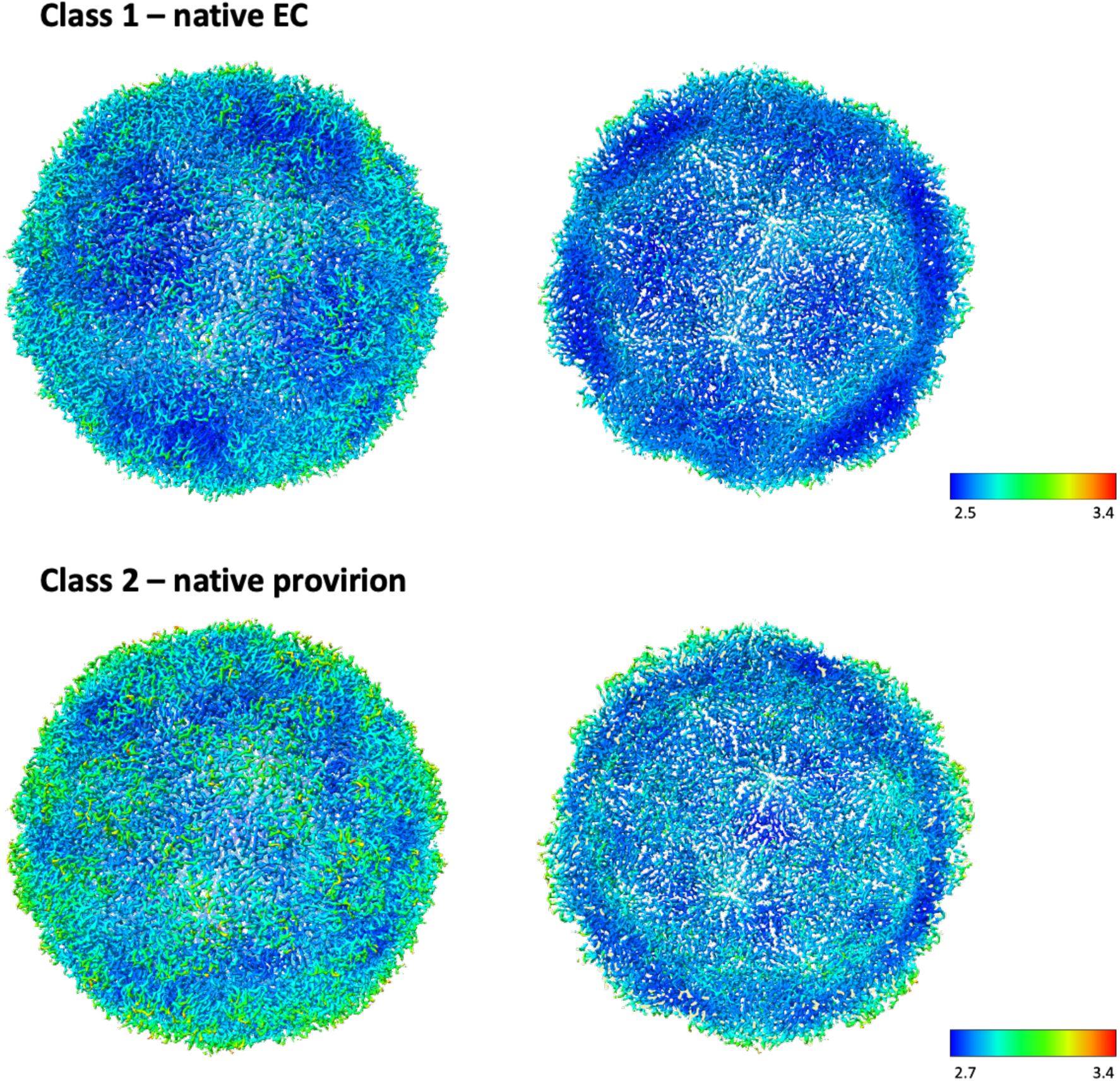
Isosurface representation of native EVA71 E096A particles: Complete (left) and sectional (right) isosurface representation of refined native EVA71 E096A particles. Particles coloured by local resolution, indicated in map-specific key. Class 1, native EC was resolved to 2.5 Å and, Class 2, the genome-containing provirion resolved to 2.7 Å. Maps displayed at 1α.

A model was built into the native EC map by rigid body fitting the 3vbs (EVA71 native virion) model using UCSF Chimera. The docked model and map were opened in Coot, the sequence corrected to match the E096A provirion sequence and residues which did not have supporting density were removed, including VP1 residues 1-58, VP4 residues 44-69 and VP2 residues 10-33 (VP0 79-102). Where the experimental density permitted, removed residues were then remodelled *de novo*, including residues from VP0 102 to 65. The model then underwent several rounds of manual adjustment in Coot and automated refinement in Phenix [45, 46].

Inspection of the final model showed a striking resemblance to the PV (PDB: 1pov) and the HRVC (PDB: 5jzg) native EC structures with α-carbon rmsd values of 1.048 and 0.827 Å, respectively [24, 28]. The resemblance was most apparent in the VP2/VP3 intra-protomer interface (internal cleft). The importance of this region was first described in relation to the PV structure [24] (Fig 5). Similar to the PV native EC structure, the VP0 cleavage boundary was modelled to the same position, where the boundary interacts with a well conserved, and putatively catalytic histidine residue at the N-terminal end of the VP2 F-strand [24] (Fig 5). Along with this conserved localisation, we also observed the same unusual φ and ψ angles around this site, with the carbonyl oxygens of the P1 and P1’ residues roughly facing the same direction, toward the VP2/VP3 intra-protomer interface. Unlike the PV and HRVC native EC structures, which resolved 13 and 8 residues at the N-terminal end of the scissile boundary, respectively, the E096A EC model was resolved for 5 residues upstream of the scissile site (Fig 5). However, while both PV and HRVC models showed discontinuous density leading to the VP0 A_1_ β-sheet, our native EC map allowed all residues between the scissile site and the VP0 A_1_ β-sheet to be modelled.

**Figure 5:**
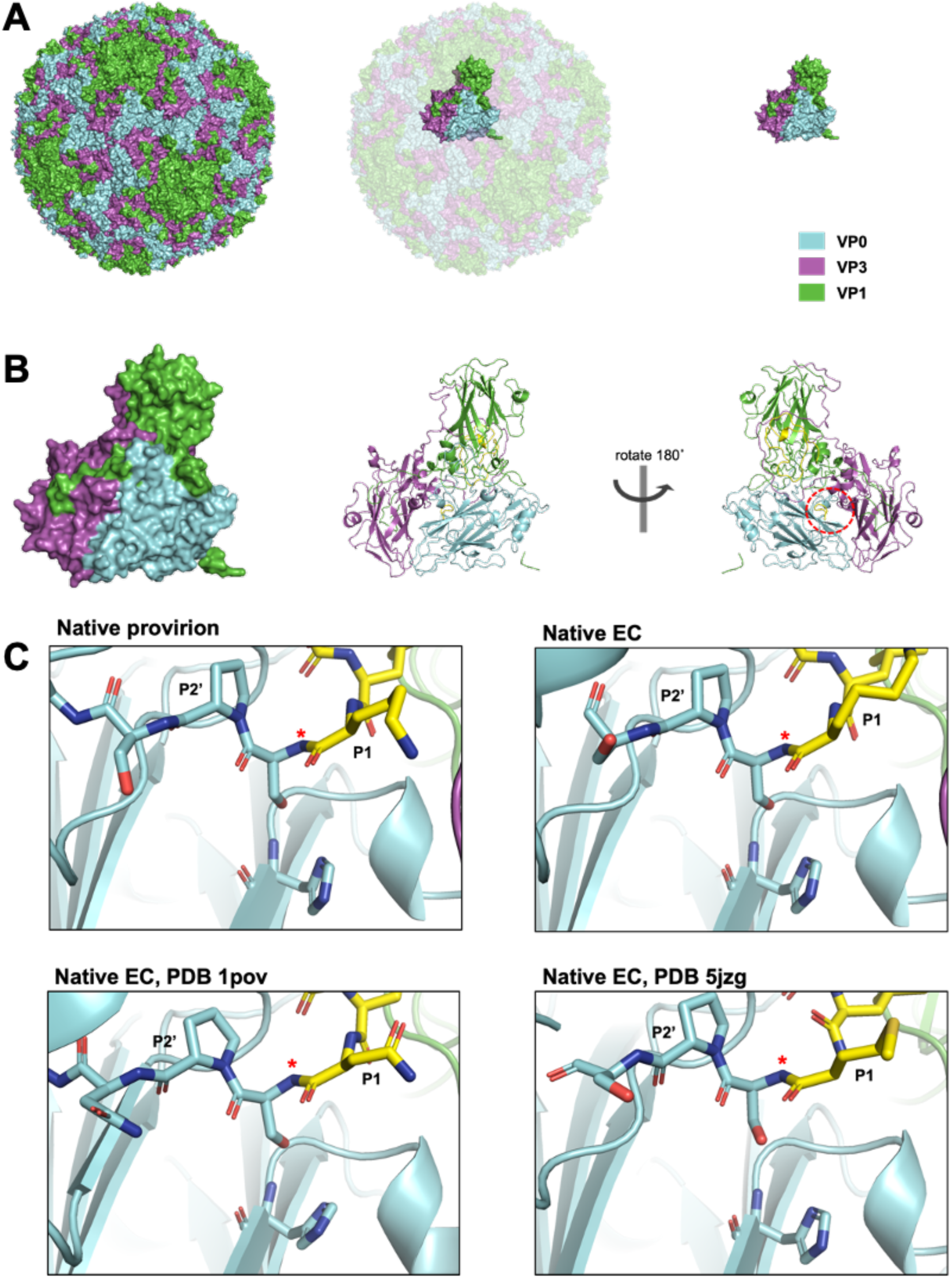
Scissile region. **A)** Surface representation of the EVA71 E096A provirion (left) with VP1 coloured green, VP0 cyan, and VP3 magenta. Position of asymmetric unit indicated in the context of full capsid (middle) and shown separately (right). **B)** Surface and cartoon representation of the EVA71 E096A protomer viewed from the external surface of the capsid, and rotated 180° along the y-axis, viewing the internal face of the capsid (right) with the approximate location of the internal cleft indicated as a red ellipse. **C)** Representation of EVA71 E096A provirion and native EC, PV1 native EC (PDB: 1pov), and HRVC native EC (PDB: 5jzg) scissile region including putative catalytic histidine. P1’ residues: E096A; K069, 1pov; N069, 5jzg; M069. Resolved residues upstream of the VP0 scissile boundary coloured yellow for clarity. P1 residue, scissile boundary (*), P2’ residue labelled. Proximal and distal regions of the models have been clipped for clarity.

The native EC model was subsequently rigid body fitted into the native provirion map in UCSF chimera [44]. Visible density was noted in the positions of residues 3-7 and 31-37 of VP1 (within 3vbs), thus these residues were added at these sites within the provirion model. Similar to the EC model, the provirion model then underwent several rounds of manual adjustment in Coot and automated refinement in Phenix [45, 46]. Unlike the EC map, the provirion map contained stretches of internal unoccupied globular density in several regions, potentially consistent with RNA density, suitable for assessment by focussed classification.

After final refinements were performed, the native EC and provirion models were compared and yielded an rmsd value of 0.220 Å, suggesting that there was minimal conformational change induced by genome packaging in the presence of the E096A mutation. The conformational similarity of the provirion model and published native ECs was also apparent with the PV (PDB: 1pov) and HRVC (PDB: 5jzg) native EC structure having rmsd values of 1.118 and 0.850 Å, respectively, when compared to the provirion model. Coxsackievirus A16 models of native EC, virus and virus-like particle were also considered in this context. The native virus-like particle (PDB: 5c8c) showed internal conformations similar to native ECs from PV and HRVC (PDB: 1pov and 5jzg) while the internal network of the native EC structure (PDB: 5c9a) shared features with both the virus-like particle and the mature virus. Since this is difficult to rationalise, the models were not included in further analysis [50].

Assessment of position-specific correlation coefficients suggests that much of the EC or provirion model fits well within either density map. A few regions of low-confidence are noted within each map/model pair, and there are several regions in which the density-fit differs between the two maps and model pairs (Fig S5). Regions of particular interest include the VP0 region positions 65-95 and the N-terminal region of VP1.

Several interesting interactions were resolved within the provirion structure leading to the VP0 A_1_ β-sheet which may provide insight into the RNA-dependence of VP0 cleavage in EVs.

W107 of VP0 is located within the flexible VP2 AB-region, and this residue (along with VP0 Y110) was predicted to interact with RNA in PV in 1989 [25]. More recently, the technological advances associated with cryoEM and asymmetric reconstruction have allowed resolution and modelling of RNA interactions with VP0 W107 in both bovine EV (BEV) (PDB: 6thd [26]) and HRV 14 (PDB: 7bg6 [27]). In each instance, structures were generated from mature virions, post VP0 processing. Curiously, within our provirion structure, rather than being involved in genome interactions, this tryptophan participates in a pi-pi stack with VP0 Y078, also forming a pi-cation stack with VP0 R081, and R081 with Y078 with distances of 5.5, 3.8, and 4.7 Å, respectively. In this conformation, there is no space available for RNA stacking interactions on this tryptophan residue (VP0 W107) (Fig 6).

**Figure 6:**
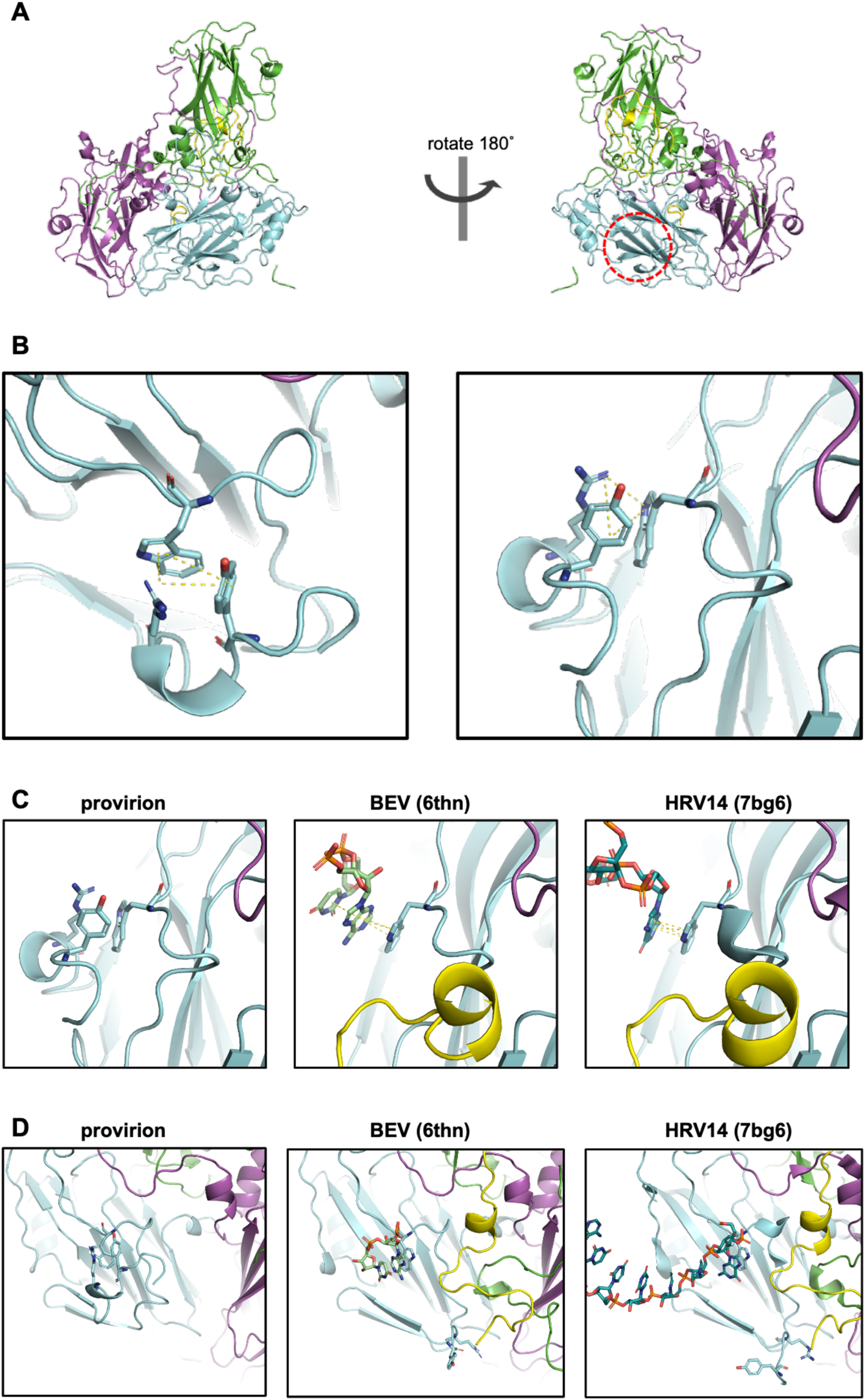
Pi-stacking residues and W107. **A)** Approximate location of the Y078, R081, W107 interaction indicated in the context of cartoon representation of E096A mutant EVA71 provirion. VP0 residues 12-69 (corresponding to VP4); yellow, residues 70-323; cyan, VP3; magenta, VP1; green. **B)** Cartoon and stick representation of W107 interacting with Y078 and R081 in EVA71 provirion (E096A), indicated pi-pi and pi-cation interactions with dotted yellow lines. **C and D)** Cartoon and stick representations of W107 interactions in EVA71 E096A provirion (left), and RNA interactions within the mature virion of BEV (6thn) and HRV 14 (7bg6), also displayed as stick Y078 and R081. VP0; cyan, VP3; magenta, VP1; green.

Globular density was noted within the pocket which sits between W107 and the VP0 paired A-sheets and to better understand the shape of this density, the map was processed through a low-pass filter at 5 Å (Fig S6). A model was built and refined by placing an RNA dinucleotide within this density, the model underwent successive cycles of manual adjustment in Coot and automated refinement in Phenix (Fig S6). The final refinements showed correlation coefficients for the dinucleotide of 0.3-0.35, thus the density fit was not satisfactory, and this region appeared inadequately resolved to allow RNA to be accurately modelled.

To better resolve the globular density observed within the provirion in proximity to W107, and at other sites, and to attempt to clarify the potentially flexible conformations seen in VP1 and VP0, focussed classification was performed. A cylindrical mask was placed covering the pertinent region within a reference asymmetric unit, and the mask part-way in the capsid shell (Fig S7). Particles were separated into 12 potential classes (regularisation parameter 30) and after 3D classification two classes contained particles, the minority class contained 36.15% of data and was resolved to 2.8 Å after final reconstruction and the majority class contained 63.85% of data which was resolved to 3.2 Å after final reconstruction, these maps were then inspected for features of interest. Both focus classes retained the additional density in proximity to VP0 W107 but the local resolution was not sufficiently improved to distinguish clear RNA structure. After low-pass filtering the maps to 5 Å additional partially ordered density was observed within the trefoil shaped depression below the scissile bond in the inner surface at the VP0/VP3 intraprotomer interface, consistent with the predicted RNA interacting moiety described in Basavappa *et al.* (1994) [24] (Fig S8). Additionally, we noted partially ordered density linked to the A_2_-B loop as well as globular density in proximity to the 2-fold axis, which appeared consistent in shape with RNA, but we cannot exclude the possibility that this density may represent some of the unresolved VP1 N-terminal residues.

### Phylogenic analysis

To assess the evolutionary consistency of the proposed mechanism of RNA-dependent maturation, detailed bioinformatic assessment was performed and phylogenic relationships assessed for VP0 cleaving picornaviruses. Of the viruses which are predicted to cleave VP0 (based upon ICTV classification), three major outgroups exist when phylogenic trees are generated using the whole viral polyprotein sequence. While limited biological data is available for the VP0 cleaving properties of many picornaviruses, it appears that cluster I contains viruses which do not require genome packaging to induce VP0 maturation, cluster II contains viruses which require genome packaging for VP0 maturation cleavage, and cluster III contains viruses which have non-standard and delayed/refractive VP0 cleavage properties (e.g. hepatoviruses [59]), although it is not clear if this is true for all members of these groupings. On the right side of the tanglegram, classification is based upon the region of interest only, an approximately 182 amino acid stretch which spans leucine just upstream of the VP0 cleavage boundary to the histidine residue in the VP0 F-strand. The three major outgroups are absolutely maintained and remarkably similar despite the region of interest using < 10% of the sequence that is present in the full polyprotein, suggesting the importance of this region of VP0 in the classification of picornaviruses (Fig 7). Additionally, the conservation of several residues in VP0 cleaving picornaviruses, and the additional conservation of VP0 Y078/E096/W107 in RNA-dependent VP0 cleaving picornaviruses supports the importance of these residues in RNA-dependent particle maturation (Fig S9, supplementary sequence files).

**Figure 7:**
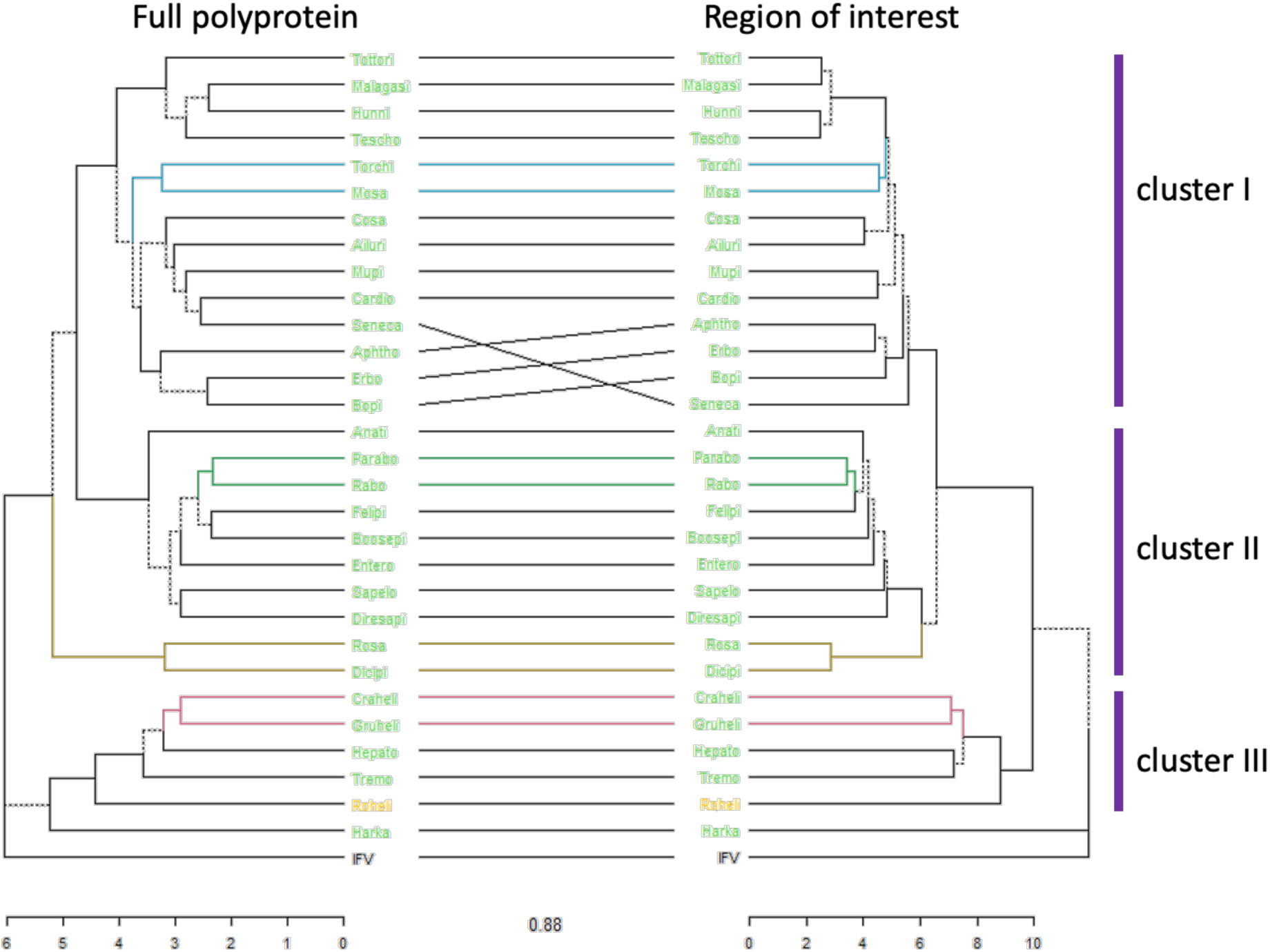
Phylogeny and tanglegram of VP0 cleaving picornaviruses: Majority consensus sequences for each ICTV-defined VP0 cleaving picornavirus were generated, and phylogenic analysis performed, consensus trees built and tanglegram constructed comparing the species-specific consensus sequence from the full polyprotein and the region of interest (spanning the leucine in the P2 position, to the histidine residue in the F-strand) which is 182 amino acids in the experimental EVA71 viral strain.

Overall, the results presented here suggest a conserved mechanism of VP0 cleavage in picornaviruses and define a mechanism for RNA-dependant maturation.

## Discussion

Through this work we aimed to shed light on the molecular details of EV assembly and maturation. There are several fundamental challenges associated with addressing these questions, not the least of which is the rapid rate at which assembling particles undergo their final maturation cleavage after genome packaging. While the functional role of a key conserved histidine residue has long been appreciated, direct evidence for the mechanism of RNA-dependence on VP0 cleavage was poorly understood [22, 24]. However, it is clear that mutation of this histidine residue is associated with inability of assembling viral particles to undergo maturation cleavage [22]. Earlier studies of the consequences of mutating this residue showed that particle assembly and genome packaging were not affected but that maturation cleavage of VP0 did not occur and the particles remained non-infectious [22].

Computational approaches revealed additional conserved sites within the viral capsid which may have roles in viral maturation. Indeed, several residues within the EV structural region are equally well conserved (>99.9%), including VP0 residue 096, which is conserved as a negatively charged amino acid. At this position glutamic acid is present in 81.5% and aspartic acid in 18.4% of sequences (7954 published sequences) representing all known EV species. Of additional interest is the relative positions of VP0 E096 in different assembly states of EV capsids. The flexibility of the VP0 A_2_B-loop, where E096 resides, suggests that it may play an important role in facilitating these conformational changes during virion assembly and maturation.

Consistent with the suggested role for E096 in facilitating conformational changes which lead to viral maturation cleavage, virus recovered from this mutant RNA did not cleave VP0, despite the presence of genome within the assembled particles (Fig 2). Consistently, infectious virus could not be recovered after serial passage (Fig S2). The precise interactions which were lacking in the E096A mutant, ultimately resulting in the accumulation of provirions, warranted further investigation.

High-resolution structures of the E096A mutant EC and provirion were generated by cryoEM (Fig 3, Fig 4) and assessment of the resultant atomic model of the EC showed remarkable consistency with previously described native EC structures. The comparison of the EC with the genome containing provirion state revealed several important features associated with this assembly intermediate. Consistent with the model proposed by Basavappa *et al.* (1994), the conserved histidine residue within the VP2 F-strand appears to interact with the VP0 scissile boundary within the E096A provirion. As in the PV EC structure, we observed stressed φ and ψ angles at the scissile boundary (Fig 5). However, unlike the PV EC model, the E096A provirion model was generated from virus particles which have packaged RNA genome. Interestingly, genome packaging in this mutant is not sufficient to induce VP0 cleavage or effectively disrupt the histidine-scissile boundary interaction. Rather, the structure of the internal region of the provirion closely resembles that of the EC. This suggests that RNA packaging in a WT virus facilitates conformational changes leading to particle maturation, but these cannot occur in the presence of the E096A mutation.

The precise mechanism of RNA induced conformational change and the RNA-dependence of VP0 maturation in EVs may be related to several stabilising interactions which centre around VP0 W107 and involve Y078 and R081. Comparing the published native EC structures, our own native EC structure and the provirion structure, several important differences in this region are apparent. In the native EC structure of PV (PDB: 1pov) the R group of VP0 Y078 and the entirety of residues 79-82 were not included because of the lack of interpretable density in the electron density maps (presumably due to disorder)[24]. In contrast in the comparable EC model of HRVC (PDB: 5jzg) Y075 (Y078 in EVA71) is modelled, along with the following 3 residues[28]. However, inspection of the density map of the HRVC model suggests that the residues in this region may have been modelled in a +1 position from residue VP0 Y075-R078 and therefore could potentially conform to the same geometry as the E096A provirion (corresponding to EVA71 residues VP0 Y078-R081) [28].

Comparisons of provirion and mature virus conformations hint at a mechanism by which the E096A mutant prevents viral maturation cleavage. In the atomic model of EVA71 provirion, the sidechain of VP0 R081 is located alongside W107 and Y078 forming stable pi-pi and pi-cation interactions (Fig 6). However, in the mature virus conformation, R081 has moved toward the VP2/VP3 internal cleft, concurrent with the elongation of the VP2 A_1_ and A_2_ β-sheets. This conformational change is not observed in the E096A mutant, and we propose that a direct interaction between E096 and R081, at least in part, facilitates this conformational shift. In addition, globular density in the region adjacent to W107, the presence of RNA forming pi-stacks with W107 in several mature EV virion structures, suggest that the displacement of R081 from its location within the assembling virion is dependent upon the packaging of genome, the interaction of genome with W107 facilitating displacement of R081, and an interaction between R081 *via* Y078 and to E096 (Fig 6).

It is worth noting that, while R081 and E096 are in proximity in several mature virus structures, the resolved sidechains are often not sufficiently close for a salt bridge to be modelled in the mature state. Additionally, within mature EVA71 virion structures W107 interacts with the VP1 N-terminal residue R017. This suggests that the interactions being described here are likely transient and final maturation in EVA71 requires additional conformational rearrangements in this region, and this includes reorganisation of the VP1 N-terminal region.

It remains unclear at which point during the internal conformational rearrangements VP0 cleavage occurs. It may be that particle maturation requires elongation of the VP2 A_1_ and A_2_ β-sheets to reposition residues upstream and downstream of the scissile site, allowing greater accessibility to the scissile boundary for the substrate (RNA side chain or cation) which ultimately facilitates cleavage at the site proposed by Basavappa *et al.* (1994) [24]. Alternatively, the RNA may mediate conformational changes, which ultimately reposition the scissile boundary away from its modelled position within this structure, to an alternative secondary active site. This second suggestion may help to explain why the released ends of the scissile boundary within mature virus structures are located a considerable distance from where they are observed in native EC structures and within our provirion structure.

Of particular interest, when this proposed model of genome dependent particle maturation was considered in the context of picornaviruses which are known to cleave VP0 in an RNA-dependent and RNA-independent manner, or not cleave VP0, several essential residues were identified.

In picornaviruses which cleave VP0 without the requirement for genome packaging (Aphthoviruses and Senecaviruses) [60–62] there are 5 conserved features:

1) Leu residue in the P2 position; 2) Glu residue at VP0 residue 74 (P5’); 3) Asp residue at VP0 position 80; 4) Arg residue at VP0 position 81; 5) and a His residue at near the N-terminal end of the VP0 F-strand.

In picornaviruses which require genome packaging to facilitate VP0 cleavage we noted an additional 3 features (Fig S9):

6) an aromatic residue at VP0 position 78 (Tyr/Phe); 7) a negatively charged residue at VP0 position 96 (Asp/Glu); 8) Trp between the A_1/2_ β-sheets.

Importantly, we have identified the coevolution of these residues/conformations and propose a functional role for each residue; the aromatic at VP0 position 78, the negatively charged residue at VP0 96, and the Trp between the A_1/2_ β-sheets (Fig S9). Whether this model is sufficient to describe VP0 maturation cleavage in all picornaviruses is unclear, due to the lack of precise molecular information about several of the lesser studied members of the family. However, in picornaviruses which do not process VP0 one or more of these 8 features are absent (or not identifiable) within the structural proteins.

The importance of this region in viral evolution is quite clearly shown in picornavirus phylogenic trees, where minimal branching differences are noted when viral relationships are defined based on either the whole viral polyprotein or the small region proposed to be critical for VP0 maturation cleavage (region of interest) (Fig 7).

It is clear that substantial conformational changes are induced by RNA packaging and that these changes are distinct from particle maturation. Indeed, employing a combined structural, bioinformatic and molecular approach facilitates the prediction of critical residues required for particle maturation. Using this approach may help to describe why in the mature virion the cleaved C- and N-terminal ends of VP4 and VP2, respectively, are located approximately 10 Å away from where they reside in the native EC and provirion structures [24]. Our analysis supports the model proposed by Basavappa *et al.* (1994), that VP2 H195 is essential for VP0 cleavage, extends evidence for the role of RNA in VP0 cleavage, and also describes a role for VP0 W107, R081, Y078 and E096 in this process. In addition, our analysis suggests an essential role for both VP0 E/D74 and D80, given the absolute conservation of these residues in VP0 cleaving picornaviruses (Fig S9). Both residues are proximal to the released ends of the VP0 cleavage boundary in mature virus structures, supporting the suggestion that they are required for VP0 cleavage, and this may utilise an aspartic protease-like mechanism. Alternatively, these residues may facilitate reorganisation of the internal protein network from the conformation seen in pre-mature states to that in mature virus. Together our data suggests a conserved mechanism for VP0 cleavage and particle maturation across picornaviruses, regardless of their dependence on genome packaging for VP0 cleavage.

Together this data suggests that although the presence of RNA is necessary for VP0 cleavage in EVs, the process of VP0 cleavage itself is not exclusively initiated by RNA packaging. The structure presented here helps to define a model for EV particle maturation which will be verified by further mutagenesis, virus characterisation and structural studies, with the aim to ultimately understand the conserved conformational changes which facilitate VP0 maturation in EVs.

## Author contributions

NJK, NJS, DJR, SJP and JMH sourced funding for this project. NJK, NJS, DJR and JMH conceived and planned experiments. NJK predicted the mutant, generated the reverse genetics system, produced, and characterised virus. NJK and EVH performed viral evolution experiments, NJK and PKH carried out qPCR based genome packaging assessment. NJK and KG prepared and purified large-scale provirion samples, JSS prepared grids and carried out cryoEM data collection. NJK processed cryoEM data with the help of JSS and TCP, NJK interpreted data with the help of JSS, DJF and JMH. NJK and JMH generated the proposed mechanism conformational stabilisation. NJK wrote the initial manuscript, and all authors were involved in editing of the manuscript and review of the data.

## Supplement

**Figure S1:**
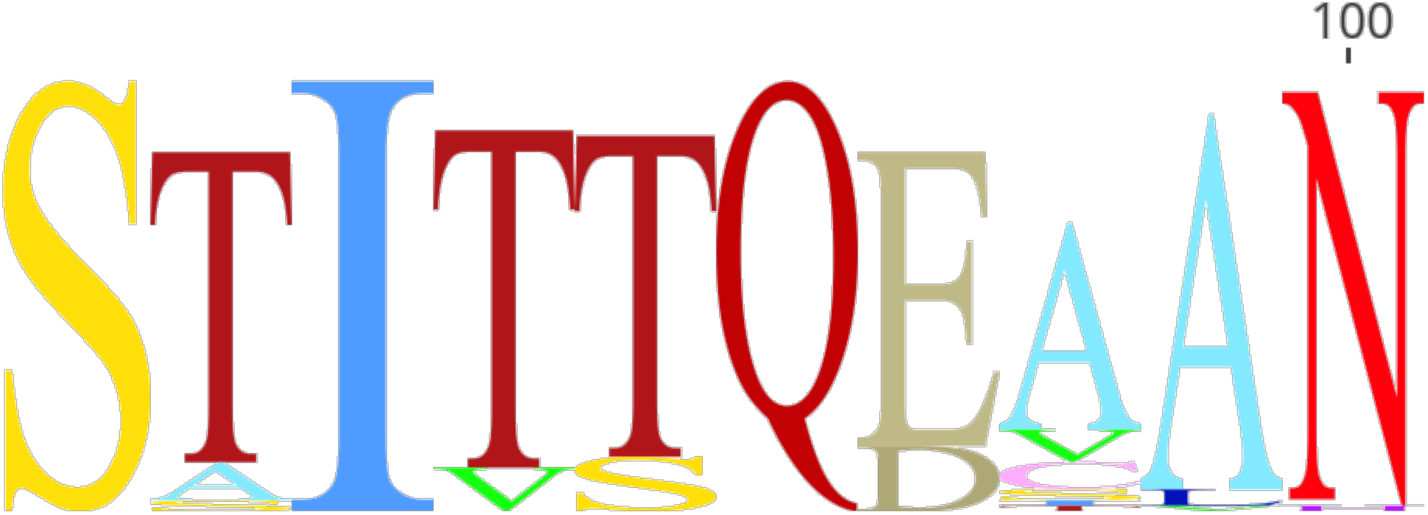
Sequence Logo of enteroviruses. Sequence logo generated from 7955 sequences, representing all enterovirus types (EV A-L, RV A-C). Displayed VP0 amino acid range 90-100.

**Figure S2:**
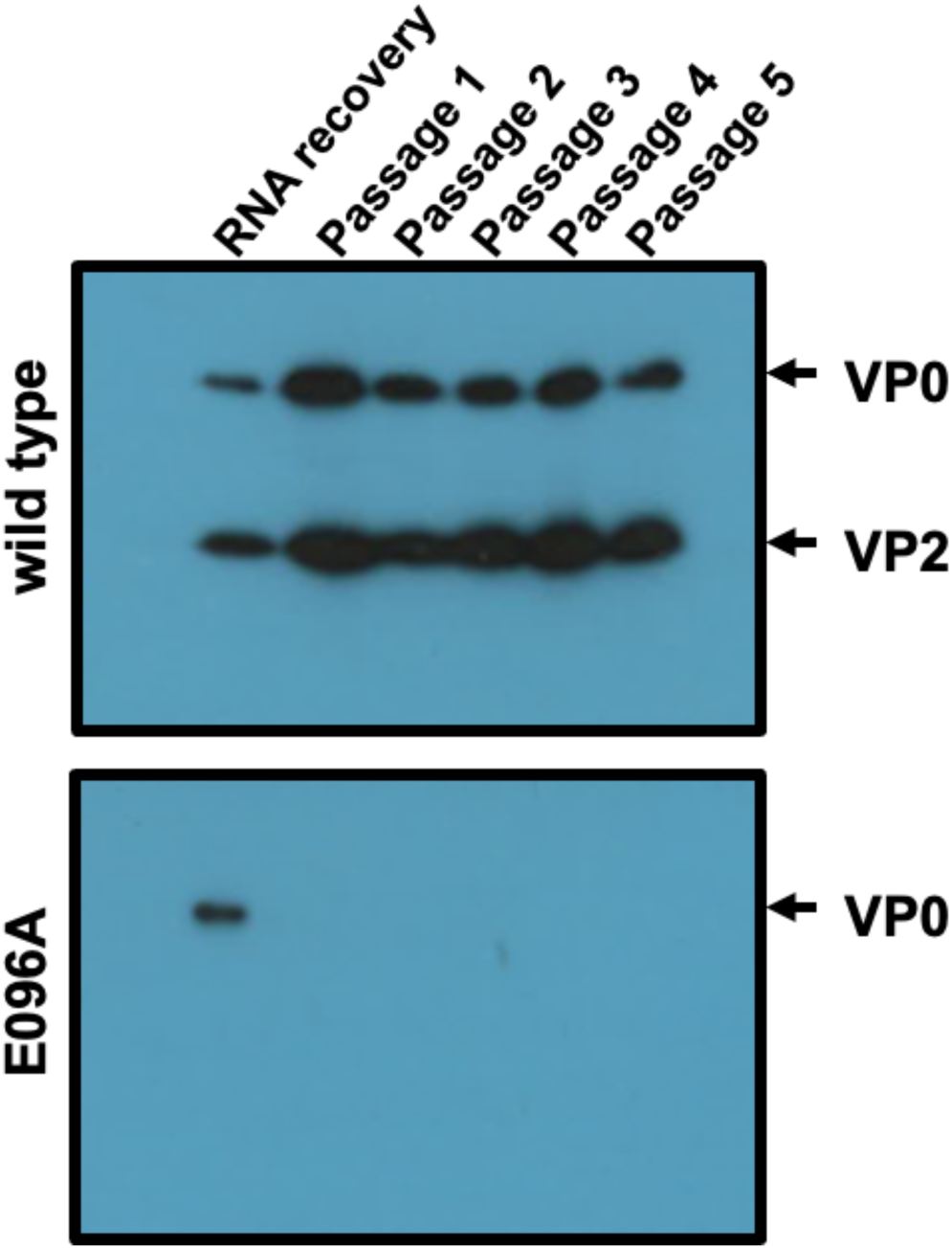
E096A mutant virus passage: Virus recovered form WT or E096A mutant viral RNA electroporated into HeLa cells. The recovered virus was passaged through HeLa cells for a total of 5 passages. No replication was detected in E096A mutant samples determined by visual inspection for signs of CPE and assessment of EVA71 proteins by western blot. Western blots show WT EVA71 and E096A mutant EVA71 samples of cell culture supernatant probed for the presence of VP0 and VP2 using mAb 979 and an anti-mouse HRP, shown representative western blot, n=3.

**Figure S3:**
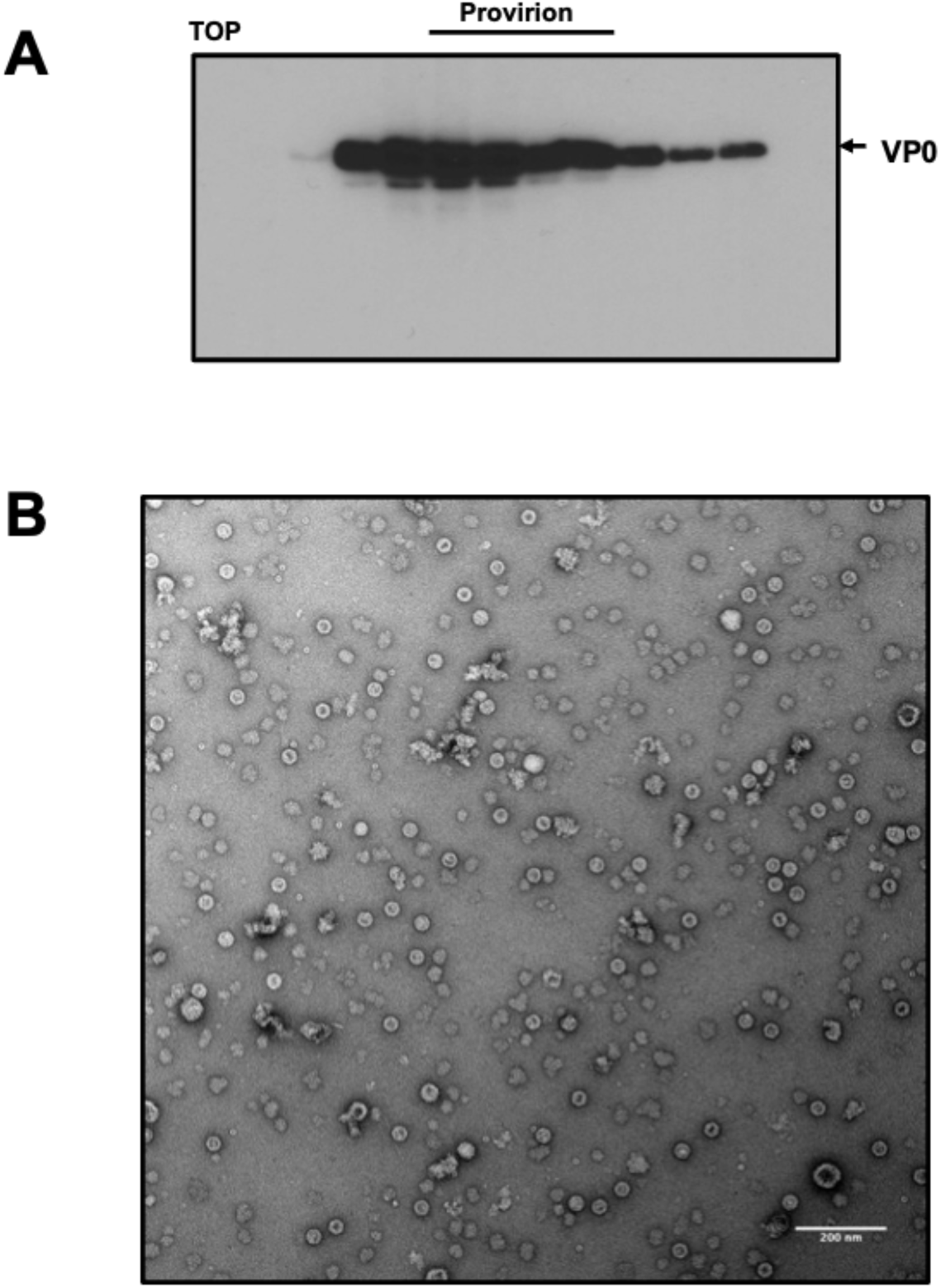
Large-scale provirion production. EVA71 E096A provirions were recovered directly from T7 transcribed RNA and purified through a 30% (w/v) sucrose cushion, before being separated along a 15-45% sucrose gradient. Fractions corresponding to provirions were subsequently diluted in PBS and underlaid with 25-45% sucrose and were further separated. **A)** Fractions were collected and assessed for the presence of VP0 and VP2 using mAb979. **B)** Peak fractions were then concentrated across a 100 kDa mwco spin concentrator with several PBS washes to remove excess sucrose. Concentrated samples were then viewed by TEM after being stained with 2% UA. Example image shown with 200 nm scale bar.

**Figure S4:**
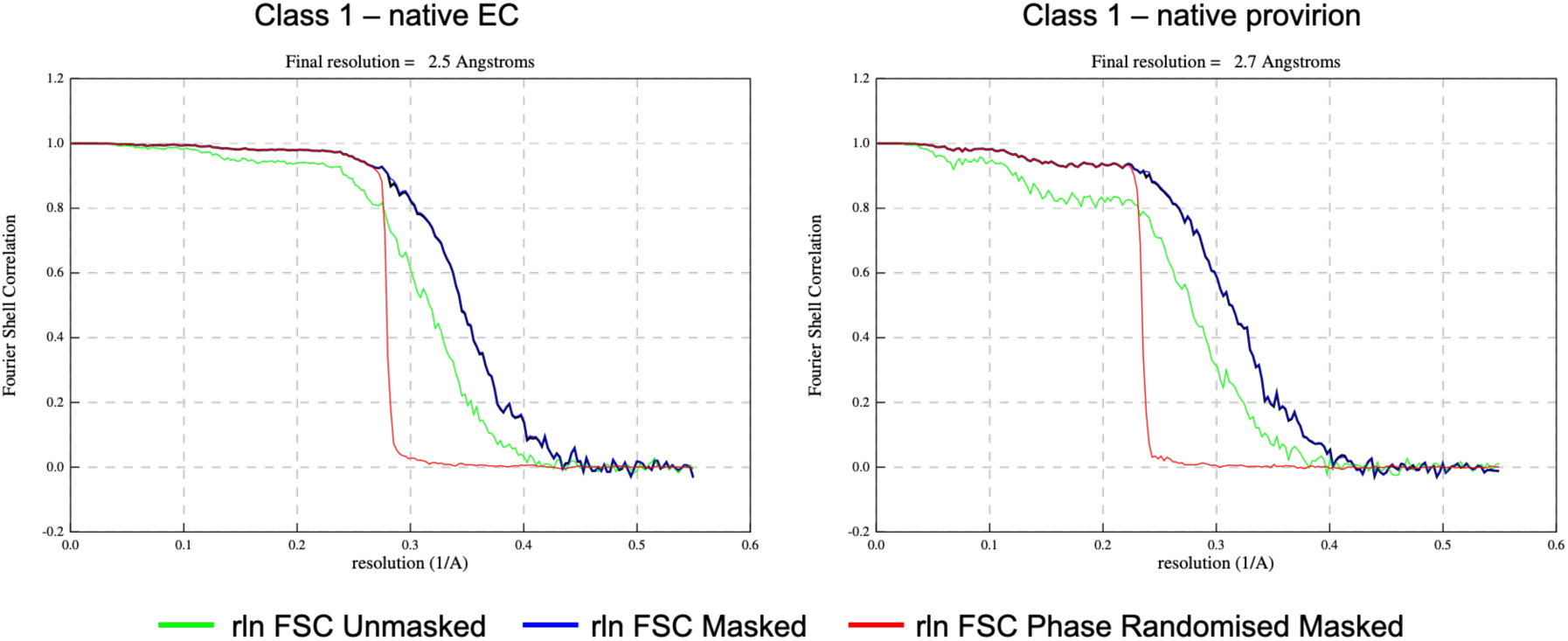
Fourier shell correlation: Fourier shell correlation (FSC) for native EC and native provirion maps, resolved to 2.5 Å and 2.7 Å, respectively, using the gold-standard (0.143).

**Fig S5:**
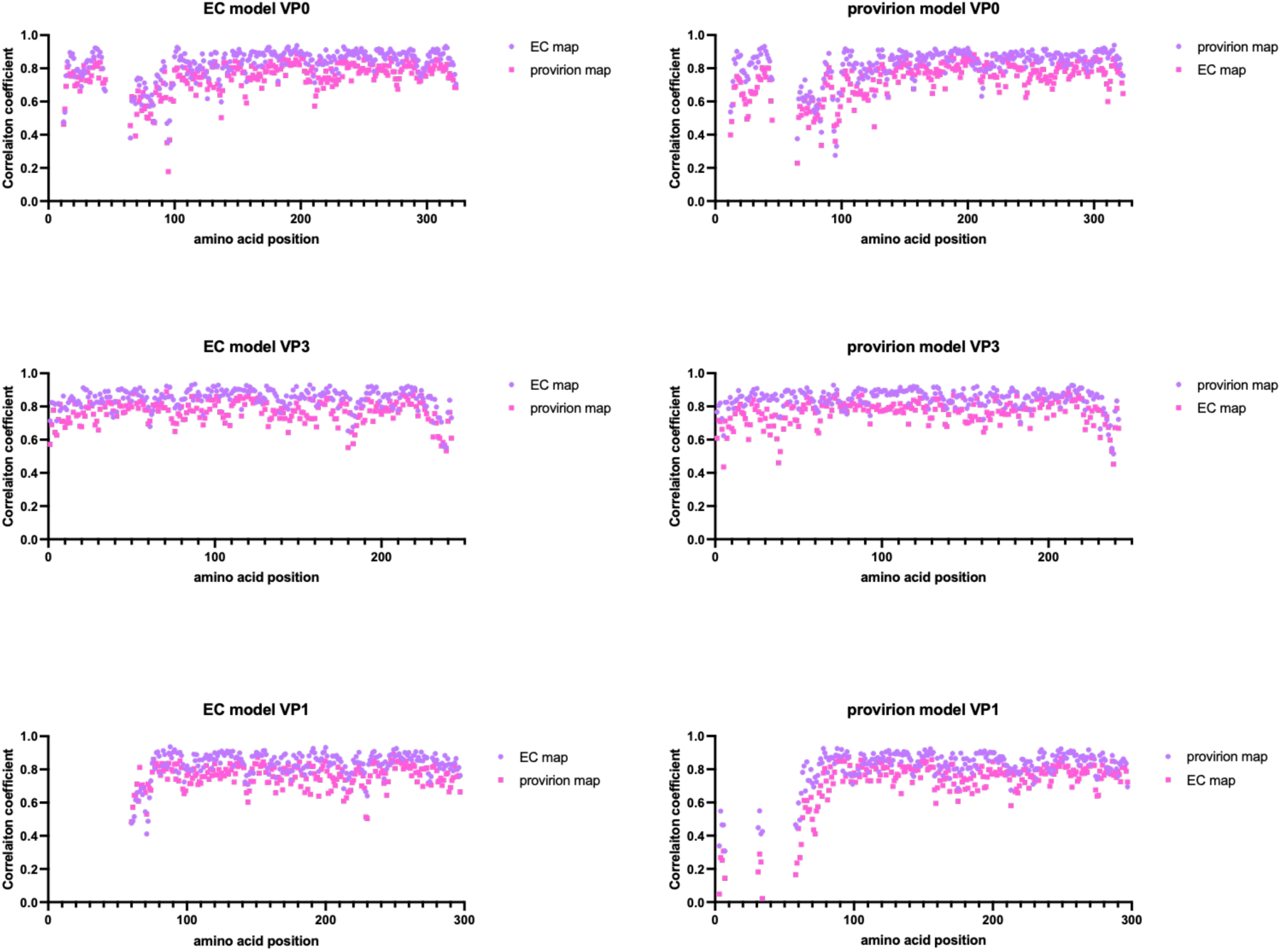
residue specific density fit: correlation coefficients for EC modelled residues (left) fitted into the EC map (purple) or provirion map (pink), and provirion modelled residues (right) fitted into the provirion (purple) or EC (pink) map. *in both instances the cognate map/model pair are displayed in purple.

**Figure S6:**
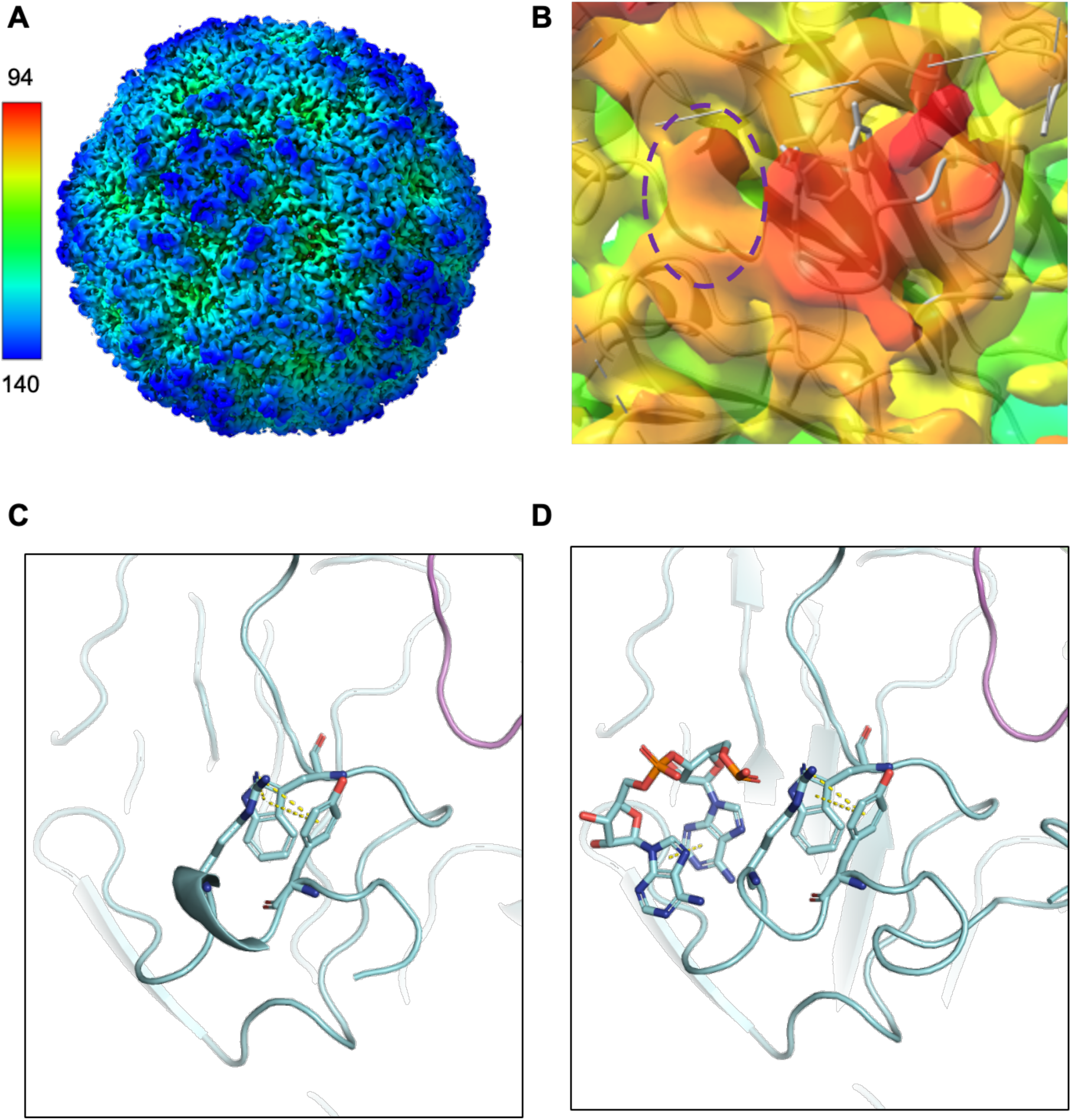
Low-pass filtered map of EVA71 E096A provirion: The EVA71 E096A provirion was processed through a low-pass filter to 5 Å. **A)** Full map after processing and **B)** regions of interest proximal to W107, R081, Y078. Additional region of globular density indicated in the dotted ellipse. Map coloured by radial distance in Å. **C)** Refined model of EVA71 E096A provirion with VP0 Y078, R081, W107 stick model and pi-interactions displayed. **D)** as **C)** but with RNA dinucleotide with pi-interactions included.

**Figure S7:**
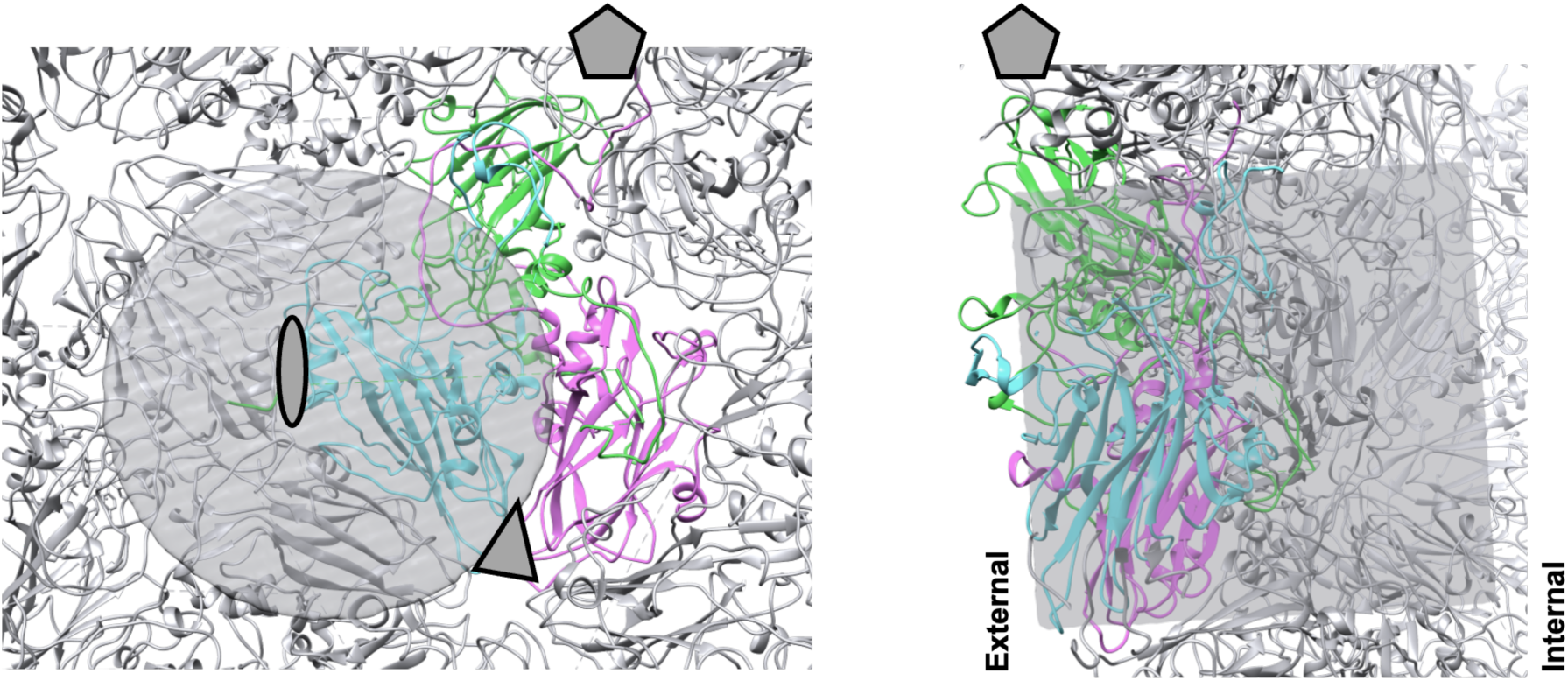
focussed classification: Location of the mask used for focussed classification. A cylindrical mask with a radius of 32 pixels and a depth of 64 pixels was placed over the reference asymmetric unit in order to cover regions with globular density and flexible regions.

**Figure S8:**
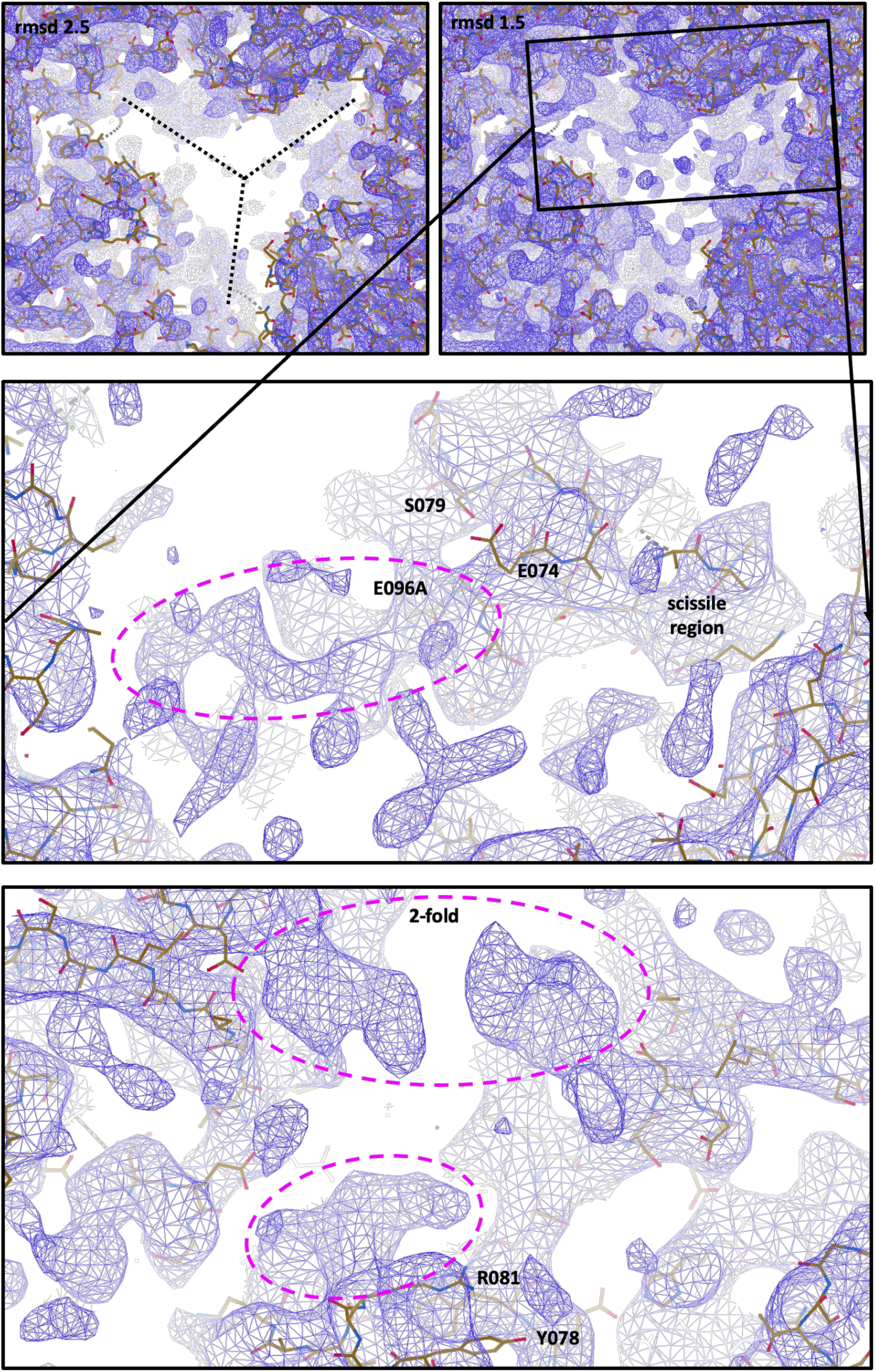
RNA organisation at the 2-fold and in the trefoil: After focussed classification was performed on the EVA71 E096A mutant provirion, the resultant map was processed through a low-pass filter. The map was displayed at 2.5 rmsd (top left) and 1.5 rmsd (top right). Density was noted in proximity to the E096A mutant residue (middle), in proximity to VP0 W107, and beneath the 2-fold symmetry axis (bottom). Regions of unoccupied density are indicated by the purple dotted ellipse.

**Figure S9:**
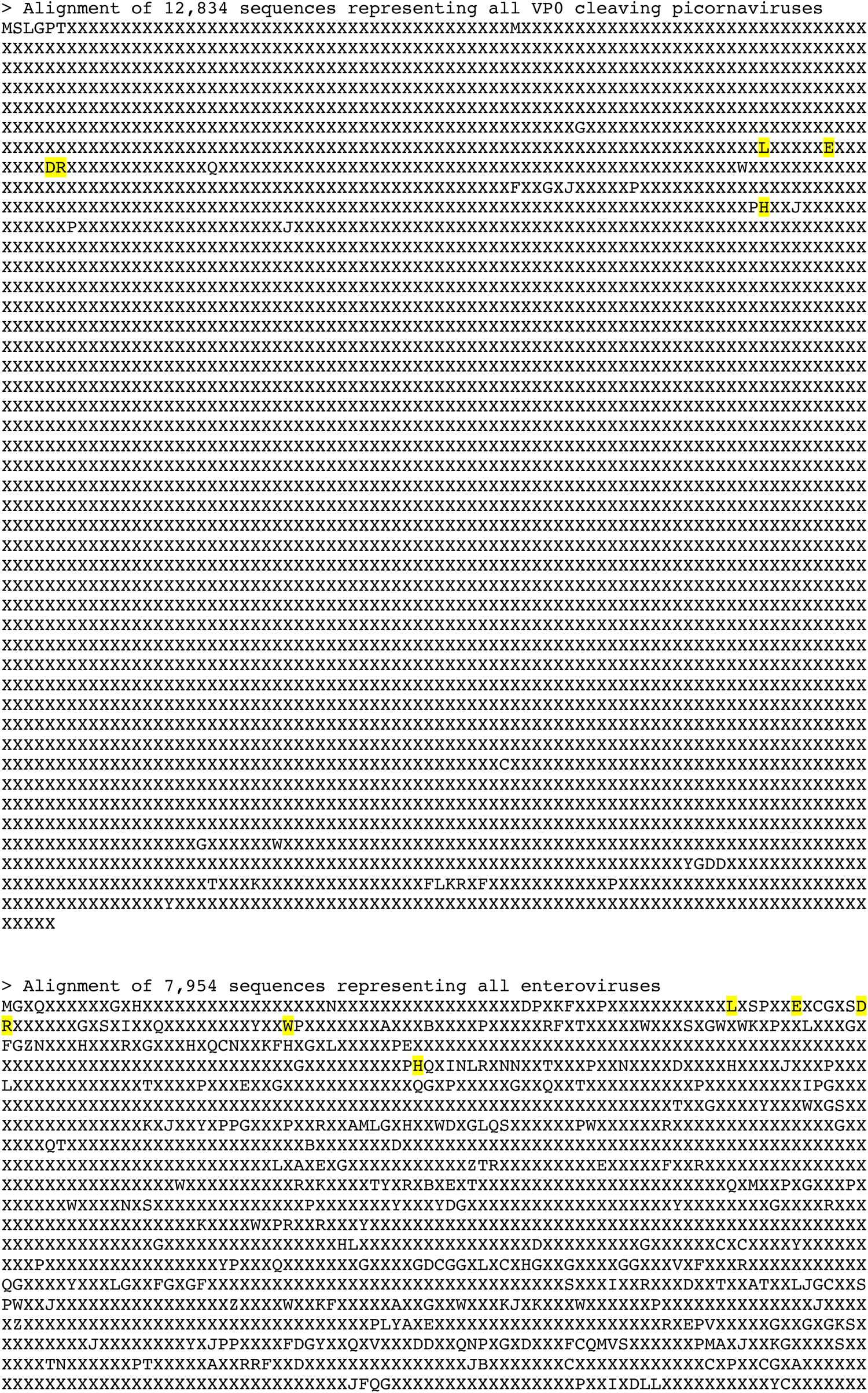
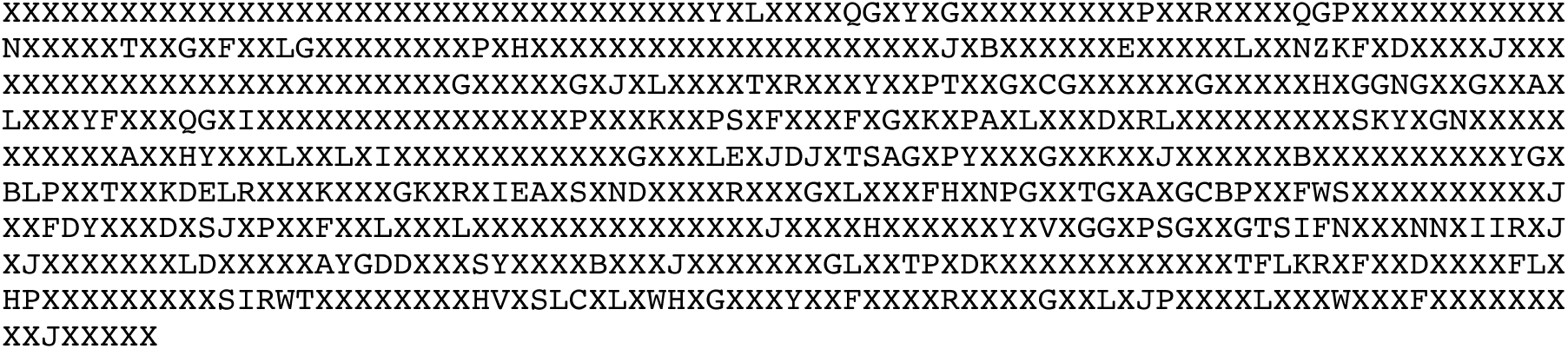
99% consensus sequence of VP0 cleaving picornaviruses: 99% consensus sequence of picornaviruses which cleave VP0 (top) and enteroviruses (bottom). Highlighted residues: **1)** a Leu residue conserved in the P2 position of the scissile boundary, **2)** the presence of a Glu residue at VP0 residue 74 (P5’), **3)** an aromatic residue at VP0 position 78 (Tyr/Phe) (does not display as a conserved residue), **4)** an Asp residue at VP0 position 80, **5)** an Arg residue at VP0 position 81, **6)** a negatively charged residue at VP0 position 96 (Asp/Glu), (does not display as a conserved residue), **7)** Trp between the A_1/2_ β-sheets, **8)** a His residue at near the N-terminal end of the VP0 F-strand.

**Table S1:** CryoEM data collection parameters.

|  | Provirion E096A |
| --- | --- |
| Microscope | FEI Titan Krios |
| Detector mode | Counting |
| Camera | Falcon IV |
| Voltage (kV) | 300 |
| Pixel size (Å) | 0.91 |
| Nominal magnification | 130,000× |
| Exposure time (s) | 5 |
| Total dose (e<sup>-</sup>/Å<sup>2</sup>) | 30 |
| Number of fractions | 41 |
| Defocus range (μm) | -0.5 to -3.1 |
| Acquisition software | Thermo Scientific EPU |

**Table S2:**
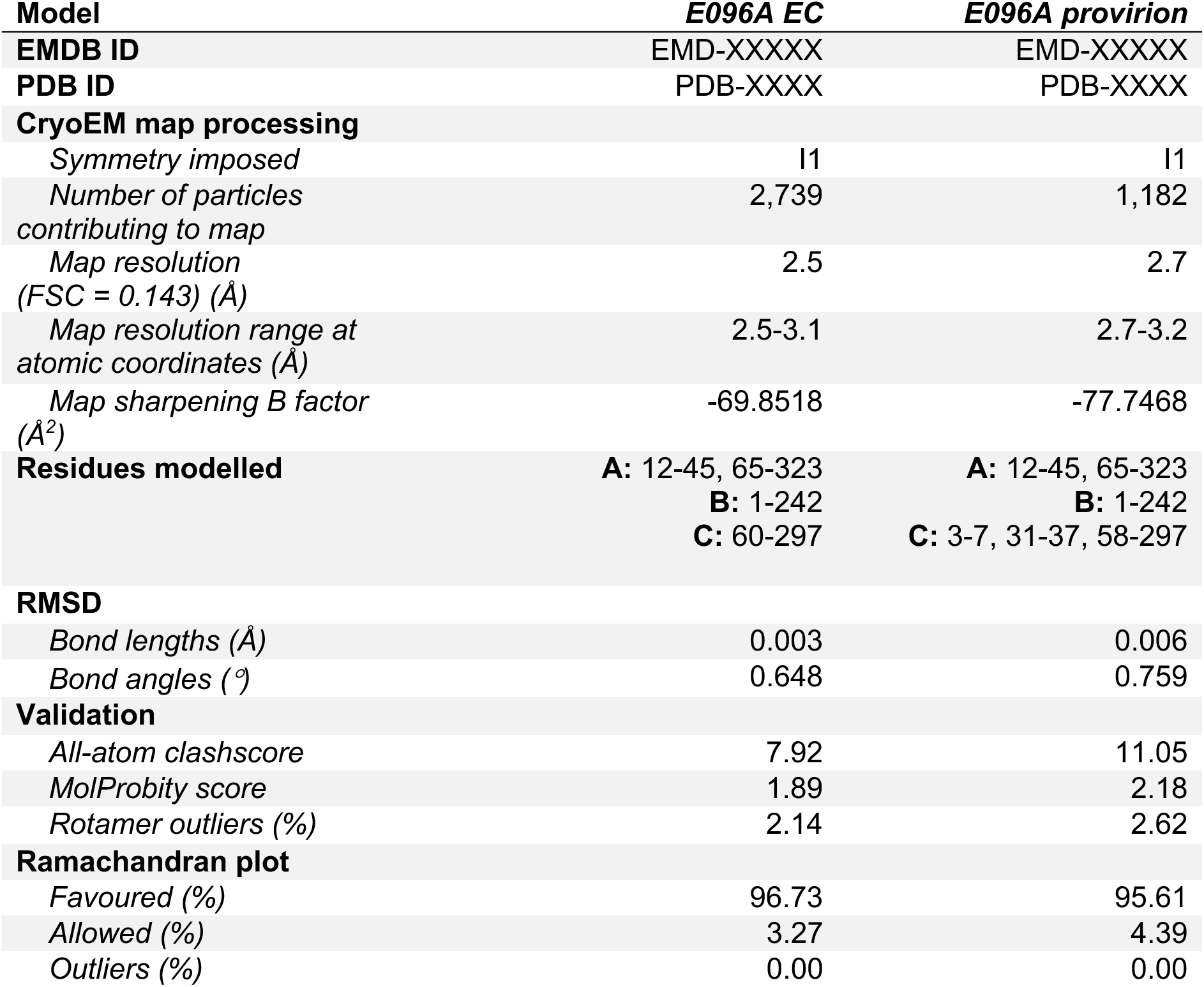
Quantitative parameters and validation statistics related to cryoEM image processing and model building.

| Model | E096A EC | E096A provirion |
| --- | --- | --- |
| **EMDB ID** | EMD-XXXXX | EMD-XXXXX |
| **PDB ID** | PDB-XXXX | PDB-XXXX |
| **CryoEM map processing** |  |  |
| Symmetry imposed | I1 | I1 |
| Number of particles contributing to map | 2,739 | 1,182 |
| Map resolution (FSC = 0.143) (Å) | 2.5 | 2.7 |
| Map resolution range at atomic coordinates (Å) | 2.5-3.1 | 2.7-3.2 |
| Map sharpening B factor (Å<sup>2</sup>) | -69.8518 | -77.7468 |
| **Residues modelled** | **A:** 12-45, 65-323<br>**B:** 1-242<br>**C:** 60-297 | **A:** 12-45, 65-323<br>**B:** 1-242<br>**C:** 3-7, 31-37, 58-297 |
| **RMSD** |  |  |
| Bond lengths (Å) | 0.003 | 0.006 |
| Bond angles (°) | 0.648 | 0.759 |
| **Validation** |  |  |
| All-atom clashscore | 7.92 | 11.05 |
| MolProbity score | 1.89 | 2.18 |
| Rotamer outliers (%) | 2.14 | 2.62 |
| **Ramachandran plot** |  |  |
| Favoured (%) | 96.73 | 95.61 |
| Allowed (%) | 3.27 | 4.39 |
| Outliers (%) | 0.00 | 0.00 |

